# Dynamic belief representation and updating through learned attractor-like dynamics in the frontal cortex

**DOI:** 10.64898/2026.09.17.752495

**Authors:** Sandra Romero Pinto, Jay Hennig, Mark Burrell, Daigo Okada, Célia Benquet, Daniel Regester, Yoh Isogai, Scott Linderman, Samuel G. Gershman, Naoshige Uchida

**Affiliations:** Department of Molecular and Cellular Biology, Harvard University, Cambridge, MA, USA; Center for Brain Science, Harvard University, Cambridge, MA, USA; Department of Psychology, Harvard University, Cambridge, MA, USA; Department of Neuroscience, Baylor College of Medicine, Houston, TX, USA; Neuroengineering Initiative, Rice University, Houston, TX, USA; Sainsbury Wellcome Centre for Neural Circuits and Behaviour, University College London, London, UK; Department of Statistics, Stanford University, Stanford, CA, USA; Wu Tsai Neurosciences Institute, Stanford University, Stanford, CA, USA

## Abstract

To act adaptively, animals must infer hidden states of the world from incomplete sensory information and update beliefs as new observations accrue. While dopamine signals are well explained by reinforcement learning models that incorporate belief states, how the brain implements belief-state inference remains unknown. Prior modeling work showed that recurrent neural networks trained to predict value (Value-RNN) develop task-specific, attractor-like dynamics that mirror evolution of beliefs. Here we performed high-density electrophysiological recordings from orbitofrontal (OFC) and other brain areas of mice performing two variants of a Pavlovian task that differ in reward probability, which produce different within-trial belief dynamics. We find that OFC population activity exhibits task-specific attractor-like dynamics that mirror the within-trial dynamics in the Value-RNN. These dynamics are absent in motor and olfactory regions, are not explained by behavioral differences, and emerge progressively with learning. Our findings indicate that the brain approximates belief-state inference through learned, task-specific attractor-like dynamics.

**Highlights:**

- High-density neuronal recording on two tasks differing in reward reliability
- OFC activity shows task-specific attractor-like dynamics mirroring belief dynamics
- These dynamics are absent in motor and olfactory areas
- Belief-like attractor dynamics in OFC emerge progressively with learning

## Introduction

Animals continuously form and update beliefs about hidden states of the world to act adaptively. Consider waiting for a train: if the schedule is reliable, as in Japan, your belief that the train will arrive grows steadily as time elapses, and you wait confidently at the platform. If the schedule is unreliable, however, as with Amtrak or during a heavy snowstorm, your belief erodes as minutes pass without a train, and you begin to doubt whether it is coming at all. This example captures a fundamental challenge in cognition: when the true state of the world is not directly observable, the brain must maintain a belief state – a time-varying probability distribution over possible states – and update it continuously as new evidence (or the lack thereof) accumulates. In the example above, this belief distribution would track the probability of being in a situation where the train arrives versus the situation where it does not. How the brain implements this computation in neural circuits remains poorly understood.

The framework of reinforcement learning (RL) provides a natural formalization of this problem. When the true state of the environment is hidden, normative theory prescribes that an agent maintains a belief state over possible states and uses it to guide value estimation and decision-making^1,2^. Previous studies have shown that dopamine signals in the brain are well explained by RL models incorporating such belief states^3–8^, suggesting that the brain computes something akin to beliefs even in simple tasks. Yet these models often require hand-crafted latent task structures to be provided and offer no mechanistic account of how belief-state computations arise from neural activity, leaving open the central question of how the brain implements belief-state inference.

A promising approach is to view neural population activity as a dynamical system^9,10^. Prior work has shown that the temporal evolution of belief states can itself be expressed as a dynamical system^2,11,12^ (see Methods). Furthermore, recurrent neural networks (RNNs) trained via temporal difference (TD) learning to predict future value (Value-RNNs) naturally develop internal dynamics that mirror belief-state dynamics, without any explicit objective to represent beliefs^8,12^. These findings raise the possibility that the brain learns task-specific recurrent dynamics that approximate the evolution of beliefs over time.

However, there are alternative explanations; upstream cortical dynamics may remain generic and task-invariant ("lazy" dynamics^13,14^), delegating value learning to plasticity in downstream circuits such as the striatum without the need to approximate beliefs. Crucially, both solutions give rise to a sufficient state representation that supports accurate value predictions^12^, making behavioral readouts alone insufficient to distinguish these possibilities.

If beliefs were to be approximated with neural dynamics, this could happen in candidate cortical regions that have previously been implicated in the representation of latent task states and beliefs. One candidate area is the medial prefrontal cortex (mPFC): acute inactivation of mPFC during task performance disrupts dopaminergic signatures of belief states without affecting core reward prediction error signals, directly implicating this region in belief computation^15^. In parallel, the orbitofrontal cortex (OFC) has been proposed to function as a cognitive map of task space^16–20^. Consistent with this idea, a recent study showed that OFC is necessary for across-trial hidden-state inference; OFC inactivation selectively impaired state inference in expert animals, and latent OFC population dynamics reflected inferred task states^21^. Together, these findings collectively point to OFC and mPFC as key substrates for belief-state inference, yet the neural mechanisms underlying time-evolving hidden-state inference or within-trial belief dynamics remain unknown.

Here, to test the hypothesis derived from our previous work^12^ that belief-updating is implemented by attractor dynamics of prefrontal cortical activity, we performed high-density electrophysiological recordings from OFC, mPFC, and other brain regions in mice performing two variants of a Pavlovian olfactory conditioning task^4^ that produce distinct within-trial belief dynamics. In one task variant (Task 1), rewards were delivered every trial, sustaining a stable belief throughout the delay period. In another variant (Task 2), reward omissions introduce ambiguity about the current task state, such that the animal’s belief of being in the reward-anticipation period rises at cue onset but decays progressively as time elapses without reward, mirroring the unreliable train scenario. Our results show that OFC, and to a lesser extent mPFC, exhibits task-specific persistent activity and fixed-point dynamics consistent with the within-trial evolution of beliefs; these signatures emerge progressively with learning, and are absent in the motor and olfactory cortices. Together, these findings support the view that prefrontal cortical circuits implement belief-state inference through learned, task-specific attractor-like dynamics.

## Results

### Task design and belief-state framework

Mice were trained in one of two versions of a Pavlovian task^4^ in which one of four odor cues (A-D) predicted a possible water reward (Figure 1A). Cue A was followed by reward after a variable delay (1.2-2.8 s, mean 2 s); Cues B and C predicted reward after fixed delays of 1.2 and 2.8 s, respectively; Cue D was never rewarded. In Task 1, rewards followed Cues A-C with 100% probability, whereas Task 2 introduced stochasticity with a 10% reward omission rate.

**Figure 1.**
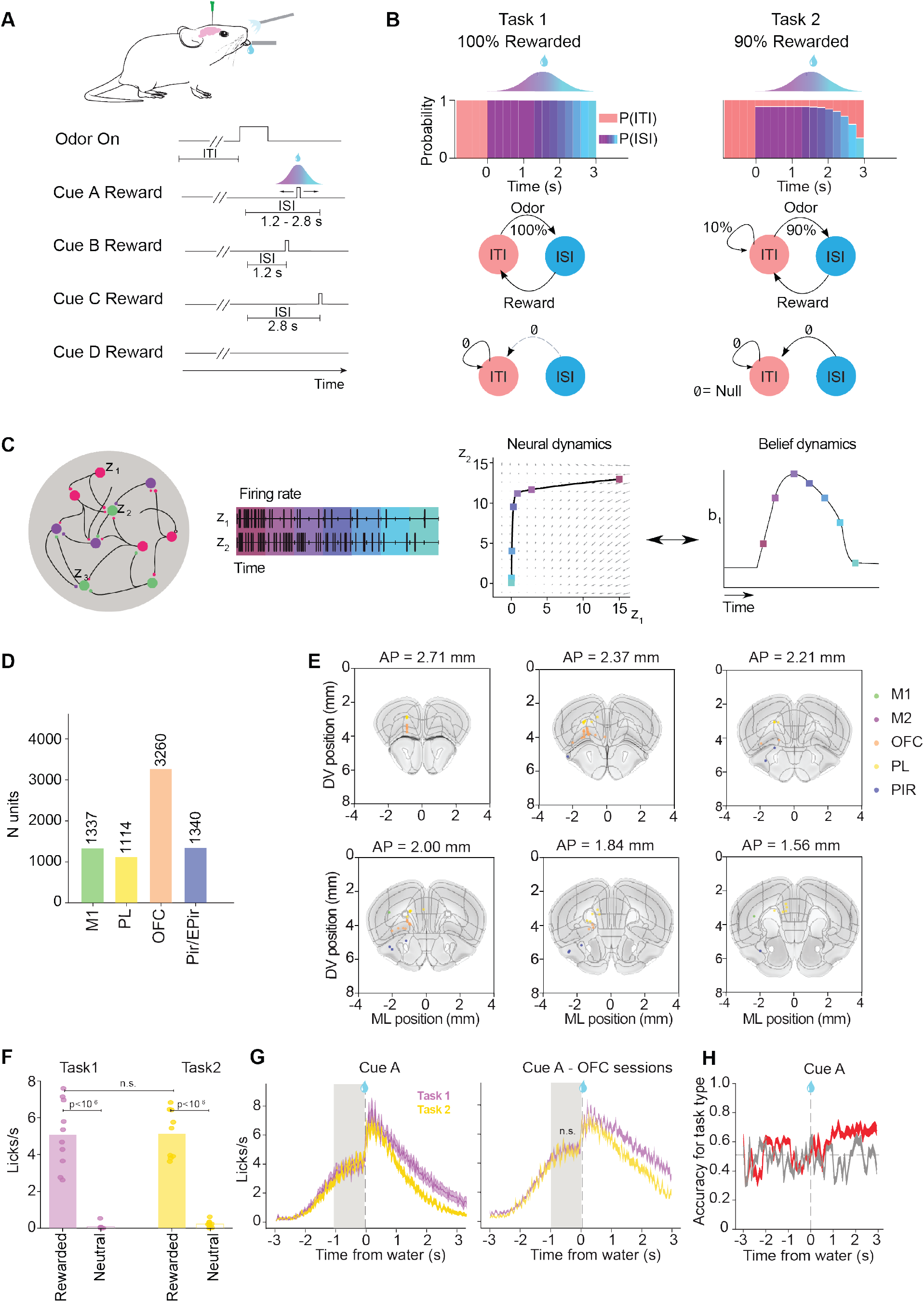
Dynamical systems framework for belief states in our task. A. Behavioral paradigm consisted of four odor cues that predicted rewards (3ul water) at different delays. Cue A predicted reward at a delay sampled from a truncated discrete gaussian distribution (1.2 to 2.8 s), Cue B predicted reward after 1.2 s, Cue C after 2.8 s and Cue D was never followed by a reward. The probability of reward delivery for Cues A-C was 100% in Task 1 and 90% in Task 2. B. Top: Probabilistic model of the belief state inference process in each task for Cue A. The model consists of two states: the ISI between odor and reward, and ITI between the reward and the next odor. The ISI state consists of 14 micro-states (200 ms bin) and the ITI of one micro-state. Middle and bottom: Transition statistics between the ISI and ITI in each task. In Task 1 each observation (cue, reward) leads to a different state (ISI and ITI respectively) while the lack of an observation (‘null’ stimulus) keeps the system in the ITI. In Task 2, the observation of a cue could lead to the ISI or keep the system in the ITI, while the lack of an observation (‘null’ stimulus) could indicate a transition from the ISI to the ITI. Thus, the states are fully observable in Task 1 but partially observable in Task 2. C. Dynamical system framework of beliefs and neural activity. In this framework, the time-varying belief state distribution is the state variable in a dynamical system that can be implemented by neural activity. D. Number of units recorded in each brain region E. Coronal views of the recording locations aligned to the Allen brain atlas. F. Average anticipatory lick rates for rewarded and neutral cues (N=10 and N=9 mice for Task 1 and 2, respectively). Effect of task type or reward for each cue type computed with a linear-mixed effects model; predictor: task type or reward type, random intercept: mouse (Effect of task type: p=0.1801, coefficient = 19.35; effect of reward for Task 1: p<10^-30^, coefficient = 102.971; effect of reward for Task 2: p<10^-20^, coefficient = 61.418). G. Left. Licking patterns for Task 1 and 2 for Cue A aligned to reward delivery averaged across sessions. Grey shaded area indicated the period used in panel D. Right. Licking patterns for Task 1 and 2 for Cue A aligned to reward delivery averaged across sessions with neural recordings in the OFC. Grey shaded area indicated the time period used in panel D (n =23, n=14 sessions in Task 1 and 2, respectively). Effect of task type computed with a linear-mixed effects model; predictor: task type or reward type, random intercept: mouse. (Effect of task type: p = 0.357, coefficient=-31.38) H. Decoding accuracy of a logistic regression model trained to predict task type with lick rates as predictors (red). Decoding accuracy for shuffled task labels (grey). Grey dotted horizontal line indicates chance level. Results are represented as mean ± SEM. M1: primary motor cortex; mPFC: medial prefrontal cortex; OFC: orbitofrontal cortex; Pir/EPir: piriform and endopiriform areas

In this task, mice could be performing belief state inference, using sensory stimuli (water and odors) to estimate which of two latent states they occupy at a given moment: the inter-stimulus interval (ISI), during which they wait for reward after an odor, and the inter-trial interval (ITI), during which they wait for the next odor (Figure 1B). In Task 1, observations fully disambiguate state transitions: the probability of being in ISI, p(ISI), jumps to 1 at cue onset and remains fixed until reward. In Task 2, however, a cue does not guarantee a transition to the ISI, so p(ISI) rises transiently after cue onset but decays over time without reward (Figure 1B).

The temporal evolution of belief states can be mathematically formulated as a dynamical system^2,11,12^, which we show can be implemented by the dynamics of neural network (Supplementary Note 1, Figure 1C). This analysis predicted one stable fixed point (i.e. an attractor) in Task 2, corresponding to the ITI belief, but potentially two stable fixed points in Task 1, where the ISI belief can be sustained indefinitely in the absence of omissions, constituting a second fixed point (Figure 1B)^12^. Consistent with this, recurrent neural networks (RNNs) trained via temporal difference (TD) learning to predict value ("Value-RNNs") developed internal dynamics that mirror these belief-state dynamics (Hennig et al., 2023; Figure 1C)^12^. Importantly, our previous study^12^ showed that an alternative model based on untrained echo-state networks^13,14^ could also support accurate value predictions by learning only downstream readout weights while keeping its dynamics fixed and identical across the two tasks. Therefore, it remains unclear whether beliefs in the brain are encoded via task-specific attractor-like dynamics, or instead whether task-specific differences in value predictions are captured by downstream plasticity changes without learning belief-like dynamics.

The two versions of the task used in our previous work^4^ provide a unique opportunity to disambiguate these two possibilities. These two task variants differ only in reward probabilities: the presence or absence of 10% reward omission trials changes the pattern of time-evolving beliefs. If an ensemble of neuronal activity encodes beliefs, it should reflect this difference. Alternatively, if there is no explicit belief state inference happening in neural dynamics, as in echo-state networks, then the activity should be similar between the two task conditions. A final possibility is that none of the brain regions we recorded from are implicated in belief computation hindering our ability to disentangle between the two possibilities.

To disambiguate between these alternatives, here we performed acute electrophysiological recordings of the activity of ensemble neurons using Neuropixels probes^22,23^. We recorded from the OFC and mPFC (prelimbic area, PL; see Methods) based on prior work suggesting these areas play a role in representing task states and beliefs (see Introduction). We also recorded from other brain regions such as the primary motor cortex (M1) and the olfactory cortices (the piriform and endopiriform areas; Pir/EPir) as controls. We recorded about 70 neurons (71.6 ± 35.5 neurons, mean ± sem) per area in each recording, resulting in 10,667 neurons in total across the 5 areas (OFC, PL, M1, Pir, EPir; Figure 1D,E).

### Anticipatory licking does not differ between tasks

Before analyzing neuronal activity, we first asked whether the two tasks differed in overt behavior. Anticipatory licking in the late ISI period, where belief dynamics diverge most, did not differ significantly between tasks (Figure 1F-G). A linear decoder trained on lick rates over time performed at chance in classifying task identity (Figure 1H), indicating that licking patterns alone could not differentiate Task 1 from Task 2. Any observed differences in neural activity across tasks, described below, are therefore unlikely to reflect overt behavioral differences.

### OFC ensemble activity mirrors task-specific belief dynamics

A key feature of Value-RNN activity is persistent deviation from the ITI baseline throughout the ISI in Task 1 but not in Task 2—reflecting sustained ISI belief in the former and its gradual decay in the latter. We examined whether ensemble activity, recorded simultaneously, exhibits this same feature using three complementary linear analyses (Figure 2).

**Figure 2.**
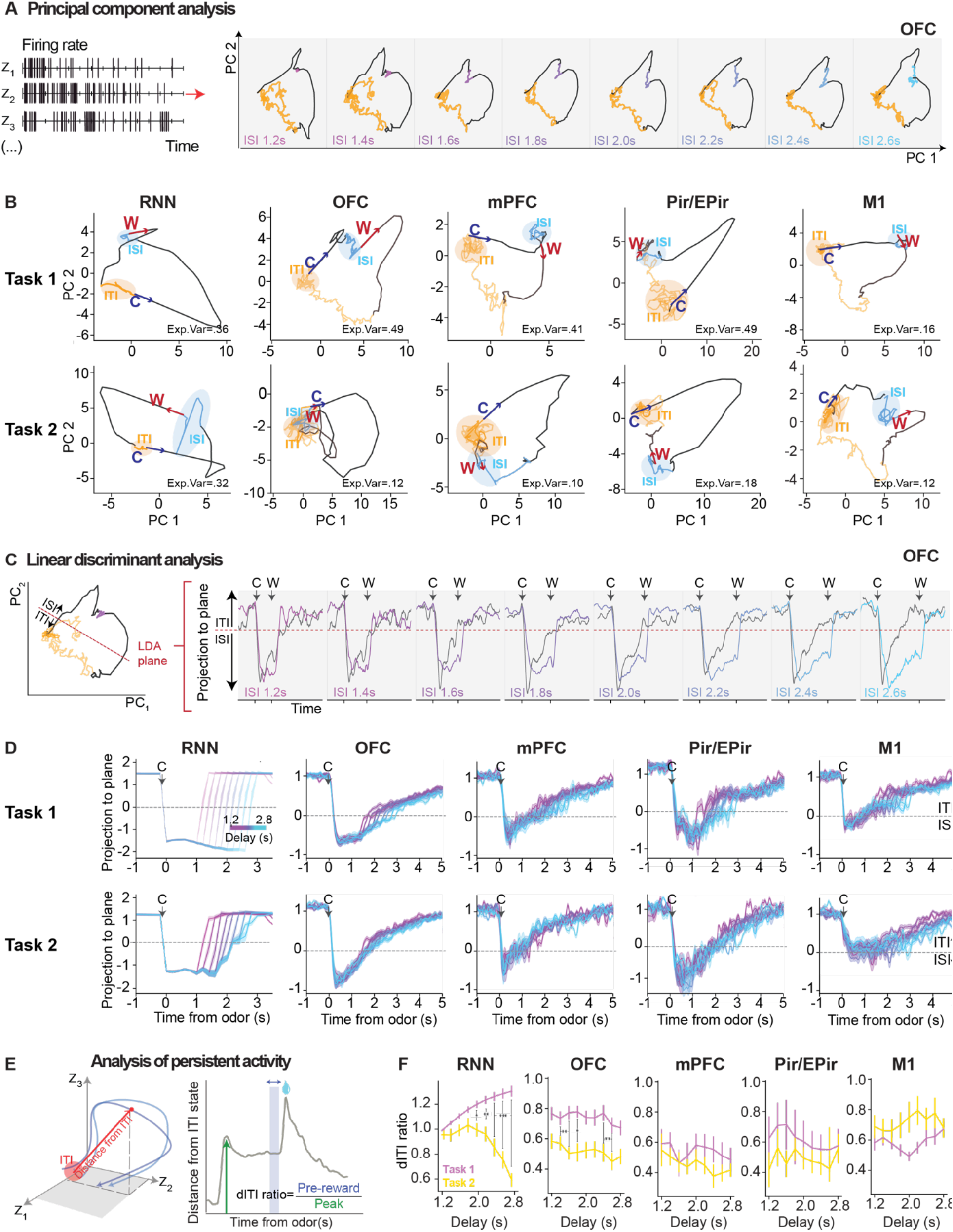
Population activity in frontal cortical areas is consistent with dynamics of beliefs and value-RNNs in Cue A. A. Schematic of principal component analysis (PCA) on the neural data for visualization. Data was projected to the space spanned by the first two principal components that maximize the variance of the data. Shown is an example session of neural activity in the OFC for Task 1, traces were averaged across trials for each ISI duration in Cue A. B. Trajectories of the first two PCA components in the recurrent neural network (RNN) activity and neural activity in the OFC, mPFC, Pir/EPir, and M1 for example sessions. Each panel shows Task 1 (top) and Task 2 (bottom) for Cue A at the delay condition of 2.6 s. The arrows show the direction of the changes in trajectory elicited by the cue (C) and reward (W). The proportion of variance explained by the first two components is indicated. C. Left: Schematic of linear discriminant analysis (LDA). Data was projected to the LDA plane that best separates the space occupied by neural activity during the ITI from the one in the ISI. Schematic shows the data in the PCA space but the full dimensionality of the data was used for this analysis. Right: example of two projected sessions for Task 1 (color) and Task 2 (grey) of neural recordings in the OFC. Values larger or lower than zero indicate that the neural activity is in the ITI or ISI side of the plane, respectively. Traces were averaged across all trials for each ISI duration for Cue A. C: cue, W: water. D. Population activity derived from the RNN and neural activity in the OFC, mPFC, olfactory areas and M1 in Task 1 (top) and Task 2 (bottom) was projected to the LDA ISI/ITI separating plane. Traces are shown for the average across all networks and all sessions for each brain region (n=22 sessions for OFC, 8 for mPFC, 7 for Pir/EPir and 16 for M1) E. Schematic of the linear analysis of persistent activity. The distance from ITI state is defined as the Euclidean distance between neural population activity at a given time bin (20 ms) and the mean activity during the ITI. Persistent activity during the ISI was quantified as the ratio of mean pre-reward distance to the peak distance evoked by the cue (i.e., maximal population deviation from baseline), termed the ‘dITI ratio.’ F. dITI ratio derived from the RNN activity and neural activity during Task 1 and Task 2 across different reward delay conditions for Cue A. Effect of task type computed with a linear-mixed effects model for each delay; dependent: dITI ratio, predictor: task type, random intercept: RNN model or mouse, corrected for multiple comparisons (n=12/12 models for Task 1 / Task 2, n=22/14 sessions for OFC, 8/7 for mPFC, 7/6 for Pir/EPir and 16/9 for M1, for Task1/Task2). See Table 1 for exact statistic values. Results are represented as mean ± SEM. ***p < 0.001; **p ≤ 0.01; *p ≤ 0.05. See also Figures S1 and S2.

**Table 1.** Effect of task on the distance-to-ITI ratio (dITI ratio) for Cue A delays.

|  |  |  |  |  |  |  |  |  |
| --- | --- | --- | --- | --- | --- | --- | --- | --- |
| **Region** | **Delay** | **Coef.** | **SE** | **z** | **p** | **p corr.** | **Sign.** | **N/task** |
| OFC | 1200 | -0.070 | 0.147 | -0.475 | 0.635 | 0.635 | n.s | 22 / 14 |
| OFC | 1400 | -0.126 | 0.090 | -1.401 | 0.161 | 0.181 | n.s | 22 / 14 |
| OFC | 1600 | -0.267 | 0.073 | -3.638 | 0.000 | 0.002 | ** | 22 / 14 |
| OFC | 1800 | -0.250 | 0.079 | -3.145 | 0.002 | 0.007 | ** | 22 / 14 |
| OFC | 2000 | -0.216 | 0.082 | -2.619 | 0.009 | 0.016 | * | 22 / 14 |
| OFC | 2200 | -0.179 | 0.098 | -1.830 | 0.067 | 0.086 | n.s | 22 / 14 |
| OFC | 2400 | -0.325 | 0.109 | -2.982 | 0.003 | 0.009 | ** | 22 / 14 |
| OFC | 2600 | -0.238 | 0.083 | -2.846 | 0.004 | 0.010 | ** | 22 / 14 |
| OFC | 2800 | -0.164 | 0.088 | -1.862 | 0.063 | 0.086 | n.s | 22 / 14 |
| mPFC | 1200 | -0.051 | 0.208 | -0.246 | 0.806 | 0.907 | n.s | 7 / 7 |
| mPFC | 1400 | -0.155 | 0.136 | -1.143 | 0.253 | 0.663 | n.s | 8 / 7 |
| mPFC | 1600 | -0.012 | 0.192 | -0.062 | 0.950 | 0.950 | n.s | 8 / 7 |
| mPFC | 1800 | -0.072 | 0.111 | -0.650 | 0.516 | 0.663 | n.s | 8 / 7 |
| mPFC | 2000 | -0.108 | 0.133 | -0.813 | 0.416 | 0.663 | n.s | 8 / 7 |
| mPFC | 2200 | -0.201 | 0.162 | -1.246 | 0.213 | 0.663 | n.s | 8 / 7 |
| mPFC | 2400 | -0.247 | 0.170 | -1.450 | 0.147 | 0.663 | n.s | 8 / 7 |
| mPFC | 2600 | -0.128 | 0.168 | -0.760 | 0.447 | 0.663 | n.s | 8 / 7 |
| mPFC | 2800 | -0.070 | 0.101 | -0.695 | 0.487 | 0.663 | n.s | 8 / 7 |
| Pir/Epir | 1200 | -0.201 | 0.213 | -0.942 | 0.346 | 0.817 | n.s | 7 / 6 |
| Pir/Epir | 1400 | -0.123 | 0.260 | -0.474 | 0.636 | 0.817 | n.s | 7 / 6 |
| Pir/Epir | 1600 | -0.257 | 0.210 | -1.222 | 0.222 | 0.817 | n.s | 7 / 6 |
| Pir/Epir | 1800 | -0.077 | 0.223 | -0.344 | 0.731 | 0.822 | n.s | 7 / 6 |
| Pir/Epir | 2000 | -0.109 | 0.172 | -0.634 | 0.526 | 0.817 | n.s | 7 / 6 |
| Pir/Epir | 2200 | -0.087 | 0.150 | -0.584 | 0.559 | 0.817 | n.s | 7 / 6 |
| Pir/Epir | 2400 | -0.091 | 0.133 | -0.687 | 0.492 | 0.817 | n.s | 7 / 6 |
| Pir/Epir | 2600 | -0.126 | 0.190 | -0.664 | 0.507 | 0.817 | n.s | 7 / 6 |
| Pir/Epir | 2800 | -0.020 | 0.175 | -0.115 | 0.908 | 0.908 | n.s | 7 / 6 |
| M1 | 1200 | -0.011 | 0.107 | -0.099 | 0.921 | 0.921 | n.s | 16 / 9 |
| M1 | 1400 | -0.050 | 0.096 | -0.519 | 0.604 | 0.921 | n.s | 16 / 9 |
| M1 | 1600 | -0.008 | 0.082 | -0.100 | 0.921 | 0.921 | n.s | 16 / 9 |
| M1 | 1800 | 0.012 | 0.068 | 0.179 | 0.858 | 0.921 | n.s | 16 / 9 |
| M1 | 2000 | 0.070 | 0.092 | 0.761 | 0.447 | 0.921 | n.s | 16 / 9 |
| M1 | 2200 | 0.034 | 0.062 | 0.549 | 0.583 | 0.921 | n.s | 16 / 9 |
| M1 | 2400 | 0.014 | 0.106 | 0.136 | 0.892 | 0.921 | n.s | 16 / 9 |
| M1 | 2600 | 0.038 | 0.128 | 0.298 | 0.766 | 0.921 | n.s | 16 / 9 |
| M1 | 2800 | -0.075 | 0.086 | -0.875 | 0.382 | 0.921 | n.s | 16 / 9 |
| Linear mixed-effects: dITI ratio ~ Task + (1\|Mouse); reference = Task 1, corrected for multiple comparisons with Benjamini-Hochberg |  |  |  |  |  |  |  |  |

We first applied principal component analysis (PCA) to visualize ensemble activity patterns across tasks. In both tasks, population activity occupied a compact, slowly evolving region during the ITI, and upon odor delivery, it was rapidly displaced from this baseline in all brain regions tested. In OFC, this displacement persisted throughout the delay in Task 1, with ISI and ITI activity remaining well-separated in PCA space. In Task 2, however, population activity progressively returned toward the ITI region as the delay extended—mirroring the decay of ISI belief (Figure 2A–B, Figure S1A-B for more example sessions). These task-dependent differences were also present for Cue C (2.8 s delay, Figure S1A-B) and were specific to OFC, mPFC and Value-RNNs, but absent in control regions: M1 activity remained persistently displaced from the ITI in both tasks, and Pir/EPir activity converged toward the ITI during the ISI in both tasks alike (Figure 2B).

Notably, the first principal component captured significantly more variance in Task 1 than Task 2, and overall neural dimensionality was higher in Task 2 (Figure S1C–D). We interpret this as reflecting the greater complexity of Task 2, in which the ambiguity of reward omissions requires the network to represent a richer and continuously evolving distribution over states^24^, rather than the discrete, stable belief maintained in Task 1. This difference in dimensionality between tasks was not followed by a greater trial-to-trial response variability in Task 2 (Figure S1E–F), supporting the view that elevated dimensionality reflects a genuine difference in the geometry of the latent space of neural activity rather than increased neural noise.

We next used linear discriminant analysis to further characterize the differences in population dynamics across tasks: We projected neural activity onto the one-dimensional, linear discriminant axis that best separates ISI from ITI activities, which allowed us to average these projections across sessions and animals. This analysis confirmed persistent ISI activity in Task 1 in both OFC and Value-RNNs, which was absent in Task 2 (Figure 2C-D; see Figure S2A-B for example single sessions). Notably, neural trajectories in Task 2 consistently diverged from those of Value-RNNs, with a systematic absence of persistent activity. This divergence was observed across all analyses and, rather than contradicting our central hypothesis, reinforces the notion of fundamentally distinct dynamical structures underlying Task 1 and Task 2.

To quantify persistence without dimensionality reduction, we computed the Euclidean distance of population activity from the mean ITI activity, normalized to the peak distance during the cue period (Figure 2E). Throughout the ISI period, the distance from the ITI activity was significantly greater in Task 1 than Task 2 in OFC (Figure 2F, Figure S2C), mirroring Value-RNN dynamics. A similar trend was observed in the mPFC albeit without reaching statistical significance and during Cue C (Figure S2D). By contrast, no task-dependent difference was observed in M1 or Pir/EPir, nor for the unrewarded Cue D (Figure S2D-E). These effects remained consistent even when accounting for the disparities in sample size across brain regions in both cues (Figure S2F). Together, these results indicate that OFC robustly exhibits task-dependent persistent dynamics consistent with belief-state encoding, with a similar but more variable trend in mPFC.

### Sustained single-neuron firing in OFC and mPFC underlies task-specific population persistence

To understand the cellular basis of task-dependent population persistence, we examined single-neuron activity. Firing rate histograms revealed sequential activation of neurons tiling elapsed time from cue onset in all regions and in both tasks (Figure 3A). However, in OFC and mPFC, many neurons in Task 1 remained active throughout the late ISI, whereas in Task 2 activity was more concentrated near cue onset.

**Figure 3.**
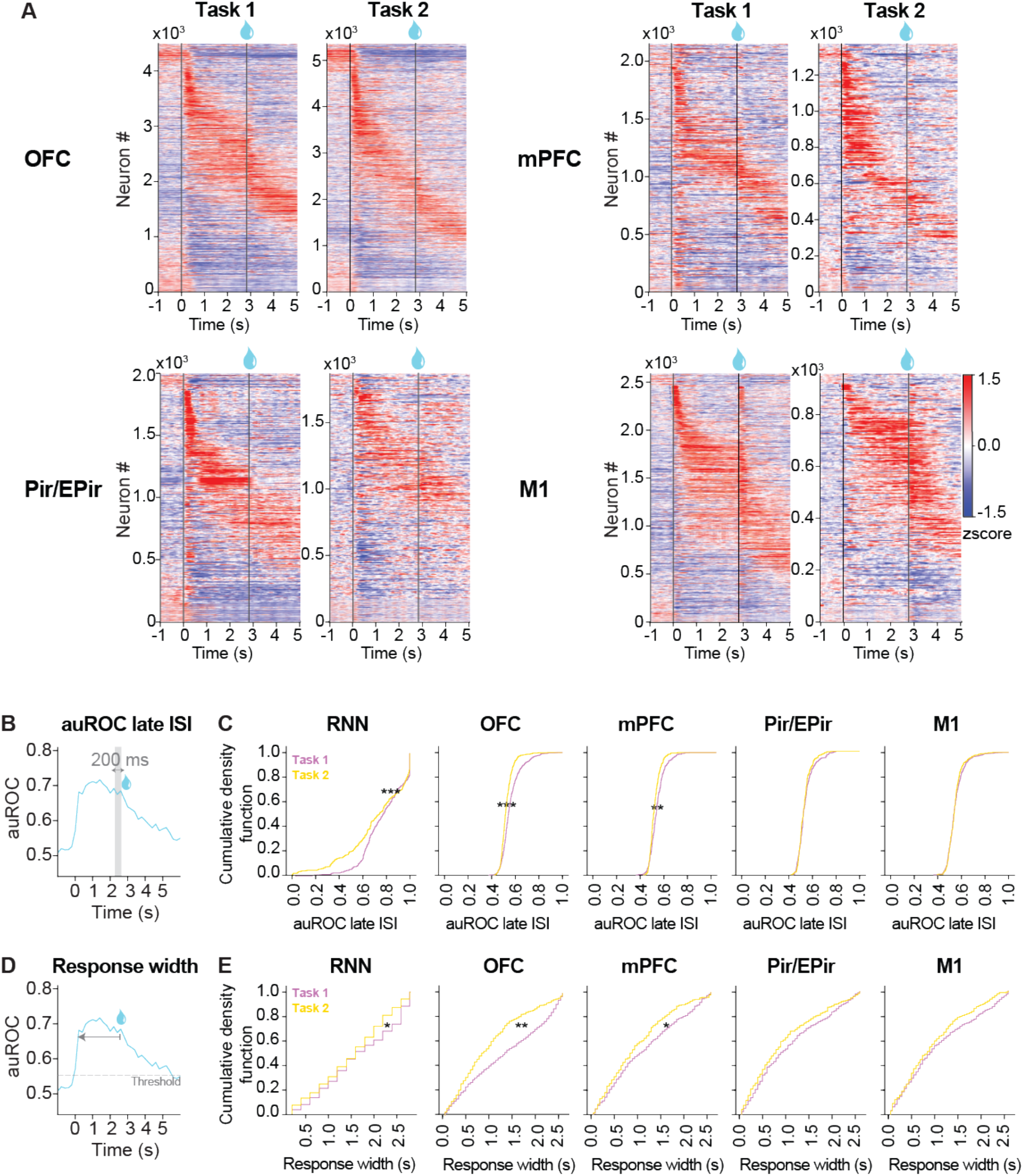
Persistent activity at the population level is supported by prolonged single neuron activity. A. Heatmaps of neural activity for a reward delay of 2.8 seconds in Task 1 and Task 2 in the OFC, mPFC, M1 and Pir/EPir. The neurons are sorted according to peak response times (PRTs) fitted on a separate half of the dataset. B. Schematic of the auROC analysis. The mean auROC was taken 200 ms before the reward period (late ISI) for each trial and averaged across trials for a given delay. C. Empirical cumulative density function (eCDF) of the auROC during the late ISI taken from Cue A trials corresponding to the two longest delays (2.6 and 2.8 s). Analysis done in the RNN activity and for each brain region. Effect of task type computed with a linear-mixed effects model for each delay; dependent: auROC during the late ISI, predictor: task type, random intercept: session (n=691/403 neurons for OFC, 313/179 for mPFC, 278/185 for Pir/EPir and 479/209 for M1, for Task1/Task2). See Table 2 for exact statistic values. D. Schematic of the response width analysis. The metric consisted of counting the time points after cue at which the auROC was above a threshold (0.55 auROC). E. Empirical cumulative density function (eCDF) of the response width based on the auROC from Cue A trials corresponding to the two longest delays (2.6 and 2.8 s). Analysis done in the RNN activity and for each brain region. Effect of task type computed with a linear-mixed effects model for each delay; dependent: response width, predictor: task type, random intercept: session (n=691/403 neurons for OFC, 313/179 for mPFC, 278/185 for Pir/EPir and 479/209 for M1, for Task1/Task2). See Table 3 for exact statistic values. ***p < 0.001; **p ≤ 0.01; *p ≤ 0.05. See also Figure S3.

**Table 2.** Effect of task on the auROC during the late ISI.

| Table 2. Effect of task on the auROC during the late ISI |  |  |  |  |  |  |
| --- | --- | --- | --- | --- | --- | --- |
| Region | Coef. | SE | z | p | Sign. | N /task |
| OFC | -0.0456 | 0.0083 | -5.5057 | 0.0000 | *** | 691 / 403 |
| mPFC | -0.0219 | 0.0099 | -2.2065 | 0.0274 | ** | 313 / 179 |
| Pir/Epir | -0.0065 | 0.0118 | -0.5512 | 0.5815 | n.s. | 278 / 185 |
| M1 | 0.0039 | 0.0076 | 0.5173 | 0.6049 | n.s. | 479 / 209 |
| Linear mixed-effects: auROC at late ISI ~ Task + (1\|Session); reference = Task 1. |  |  |  |  |  |  |

**Table 3.** Effect of task on the response width based on auROC.

| Table 3. Effect of task on the response width based on auROC |  |  |  |  |  |  |
| --- | --- | --- | --- | --- | --- | --- |
| Region | Coef. | SE | z | p | Sign. | N /task |
| OFC | -542.8098 | 134.2554 | -4.0431 | 0.0001 | *** | 691 / 403 |
| mPFC | -342.3951 | 141.3487 | -2.4223 | 0.0154 | ** | 313 / 179 |
| Pir/Epir | -179.0706 | 260.6124 | -0.6871 | 0.4920 | n.s. | 278 / 185 |
| M1 | -173.7443 | 137.4482 | -1.2641 | 0.2062 | n.s. | 391 / 166 |
| Linear mixed-effects: Response width ~ Task + (1\|Session); reference = Task 1. |  |  |  |  |  |  |

To quantify this difference, we applied receiver operating characteristic (ROC) analysis^25^, where the area under the curve (auROC) measures the reliability of firing rate deviations from baseline across trials, with values above or below 0.5 indicating excitation or inhibition, respectively. During the late ISI (200 ms before reward onset), auROC values were overall shifted toward higher values in Task 1 relative to Task 2 in OFC, mPFC and Value-RNNs (Figure 3B-C). We further computed ‘response width’, the cumulative duration for which a neuron’s auROC exceeded 0.55 or fell below 0.45 for excited and inhibited neurons, respectively, as a measure of sustained single-neuron activity (Figure 3D). Response widths were systematically broader in Task 1 than Task 2 in OFC, mPFC, and Value-RNNs (Figure 3E, Figure S3A), indicating that neurons in these regions maintained prolonged deviations from baseline throughout the ISI specifically in Task 1. This was observed in both neurons excited and inhibited by the ISI period during Cue A (Figure S3A) and Cue C trials (Figure S3B) but not in Cue D trials (Figure S3C).

### State decoding from OFC and mPFC tracks belief uncertainty

If neural activity encodes belief states—i.e., probability estimates over ground-truth task states—then those task states should be decodable from neural activity. Moreover, the structure of the uncertainty of a decoder trained to classify ground-truth states should reflect belief uncertainty. In Task 2, where belief in the ISI diminishes over time, the decoder’s estimated probability of being in the ISI should likewise decline, whereas in Task 1 it should remain stable. To test this, we trained a linear multinomial decoder to classify 14 ISI micro-states (200 ms windows) and one ITI state from population activity (Methods, Figure 4A).

**Figure 4.**
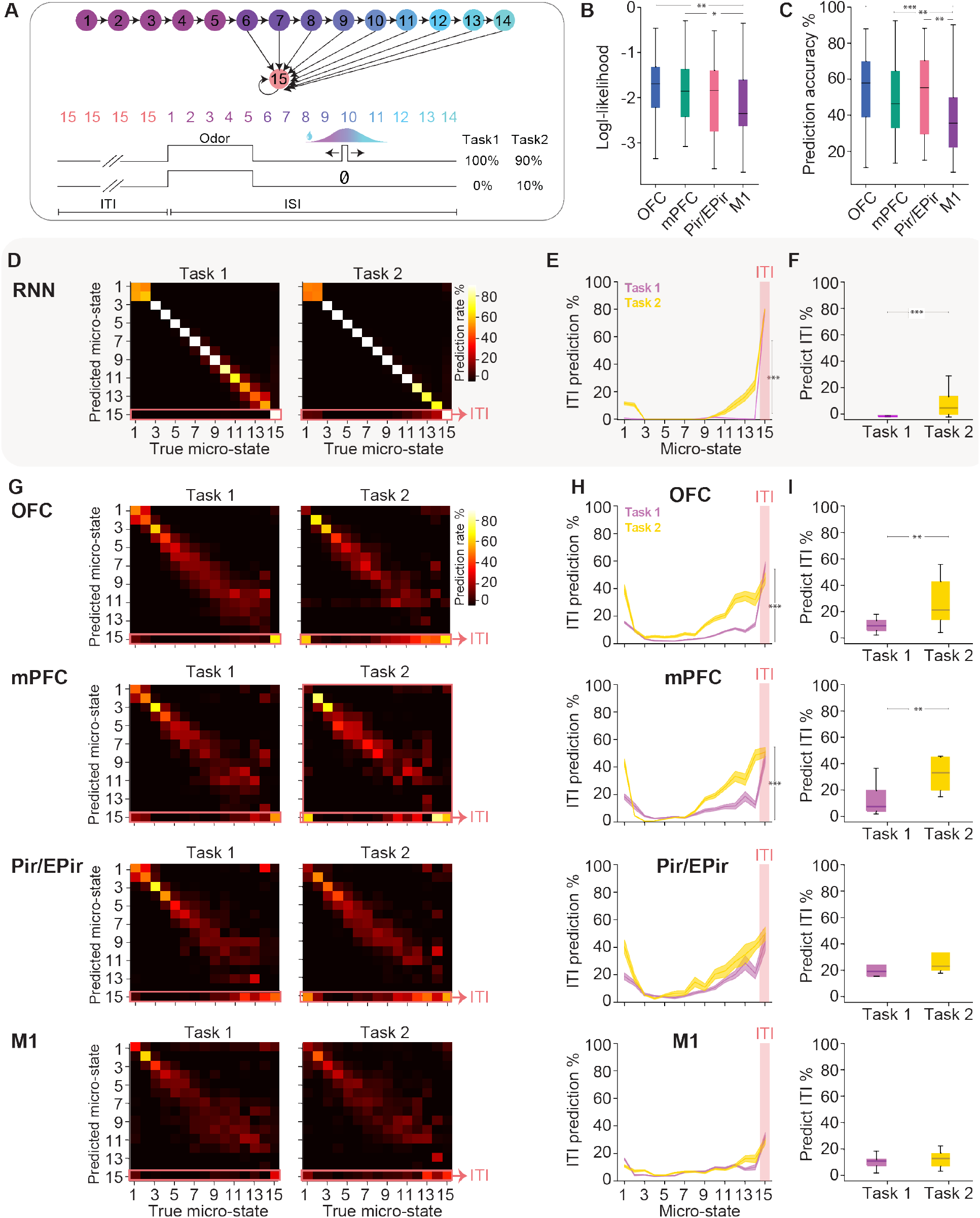
Accuracy in linearly decoding task states is consistent with neural activity belief states. A. Microstates in the belief state model. Microstates 1-14 correspond to the ISI, while the ITI consists of the single substate 15. An odor input transitions the system from the ITI microstate 15 to the first ISI microstate. In Task 1, water delivery transitions the system from an ISI microstate to the ITI with 100% probability. In Task 2, there is a 10% probability of transitioning from the last ISI microstate to the ITI in absence of water. B. Mean log-likelihood of the state decoder trained on neural activity from each brain region averaged across tasks. Effect of brain region was computed with a permutation test; pairwise comparisons were Bonferroni corrected; p =0.041 for difference between OFC and Pir/EPir, p =0.00047 for difference between OFC and M1, p=0.0027 for difference between mPFC and M1 (n= 29 sessions for OFC, 11 for mPFC, 9 for Pir/EPir and 24 for M1 pooled across both tasks). See Table 4 for exact statistic values. C. Mean prediction accuracy of the state decoder trained on neural activity from each brain region averaged across tasks; p = 0.00024 for difference between OFC and M1 and p=0.00035 for difference between mPFC and M1, p=0.0116 for difference between Pir/EPir and M1, (n= 29 sessions for OFC, 11 for mPFC, 9 for Pir/EPir and 24 for M1 pooled across both tasks). See Table 5 for exact statistic values. D. Confusion matrix of micro-state predictions from RNN activity trained on Task 1 and 2 E. Rate of ITI prediction of state decoders based on RNN activity trained on Task 1 and 2 for each micro-state. Interaction of task type and micro-state was computed with a linear-mixed effects model; dependent: ITI prediction rate, predictor: task type and state, random intercept: session. F. Rate of ITI prediction across the micro-states spanning 2.0 to 2.8 s. Effect of task type was computed with a linear-mixed effects model; dependent: ITI prediction rate, predictor: task type, random intercept: mouse G. Same as D but the state decoders were trained on neural activity from the OFC, mPFC, Pir/EPir and M1. (N=20/9 sessions for OFC, 7/4 for mPFC, 5/4 for Pir/EPir and 16/8 for M1 for Task/Task2). H. Same as E but the state decoders were trained on neural activity from the OFC, mPFC, Pir/EPir and M1. Same sample size as panel E. Interaction effect of task type and micro-state was computed with a linear-mixed effects model; dependent: ITI prediction rate, predictor: task type and state, random intercept: mouse; p<10^-10^ for OFC, p=0.00034 for mPFC, p=0.512 for Pir/EPir, p = 0.011 for M1. See Table 6 for exact statistic values. I. Sames as F but the state decoders were trained on neural activity from the OFC, mPFC, Pir/EPir and M1. Same sample size as panel E. Effect of task type was computed with a linear-mixed effects model; dependent: ITI prediction rate during the late ISI (2.0 to 2.8 s), predictor: task type, random intercept: mouse; p=0.00019 for OFC, p=0.0087 for mPFC, p=0.448 for Pir/EPir, p = 0.055 for M1. See Table 7 for exact statistic values. Results are represented as mean ± SEM. ***p < 0.001; **p ≤ 0.01; *p ≤ 0.05.

**Table 4.** Inter-region differences in the log-likelihood of a micro-state decoder.

|  |  |  |  |
| --- | --- | --- | --- |
| Region 1 | Region 2 | p-value corrected | Sign. |
| M1 | OFC | 0.0029 | ** |
| M1 | Pir/Epir | 0.4774 | n.s. |
| M1 | mPFC | 0.0518 | * |
| OFC | Pir/Epir | 0.2834 | n.s. |
| OFC | mPFC | 0.5063 | n.s. |
| Pir/Epir | mPFC | 0.4774 | n.s. |
| Permutation test corrected for multiple comparisons with Benjamini-Hochberg |  |  |  |

**Table 5.** Inter-region differences in the label accuracy of a micro-state decoder.

|  |  |  |  |
| --- | --- | --- | --- |
| Region 1 | Region 2 | p-value corrected | Sign. |
| M1 | OFC | 0.0002 | *** |
| M1 | Pir/Epir | 0.0090 | ** |
| M1 | mPFC | 0.0073 | ** |
| OFC | Pir/Epir | 0.4558 | n.s. |
| OFC | mPFC | 0.2500 | n.s. |
| Pir/Epir | mPFC | 0.8007 | n.s. |
| Permutation test corrected for multiple comparisons with Benjamini-Hochberg |  |  |  |

**Table 6.**
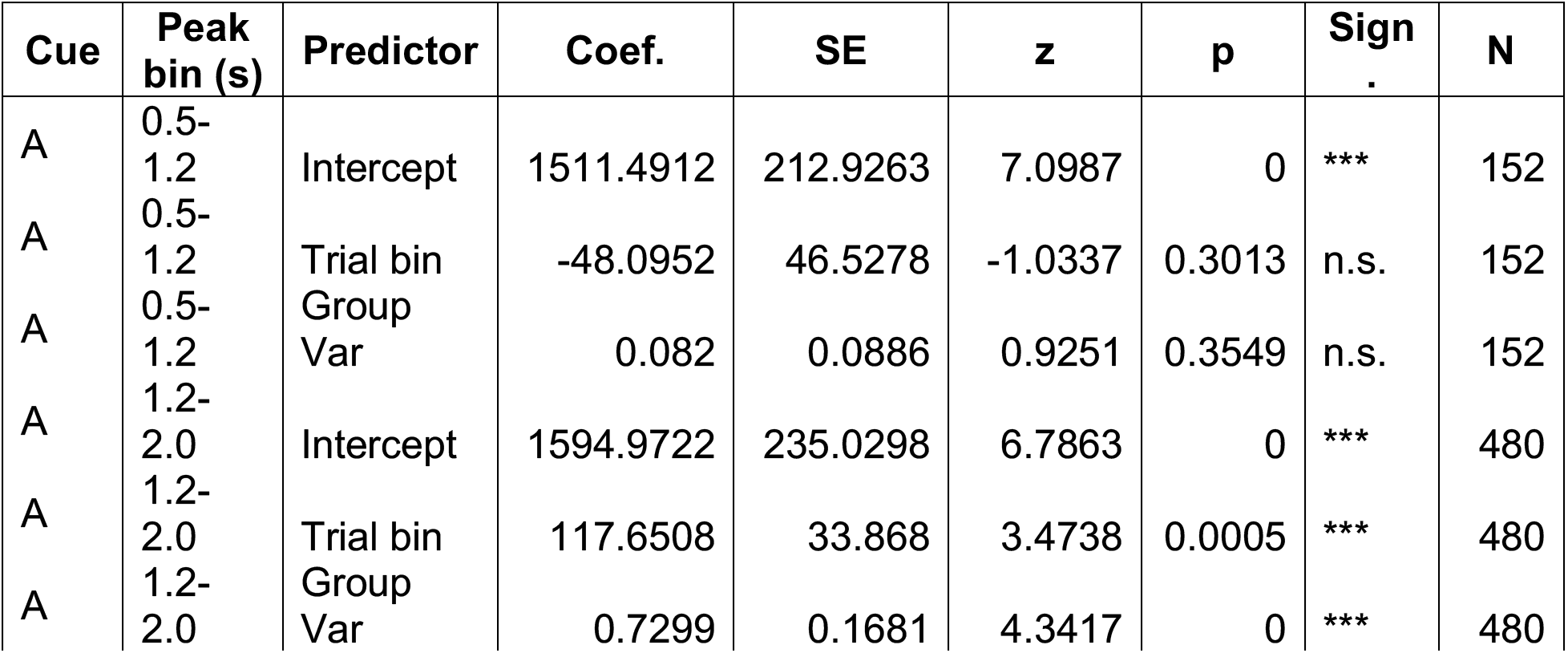
Effect of task and micro-state on the percentage of ITI prediction.

**Table 7.** Effect of task on the percentage of ITI prediction during the late ISI.

|  |  |  |  |  |  |  |
| --- | --- | --- | --- | --- | --- | --- |
| **Region** | **Coef.** | **SE** | **z** | **p** | **Sign.** | **N /task** |
| OFC | 0.1480 | 0.0633 | 2.3384 | 0.0194 | ** | 20 / 11 |
| mPFC | 0.1893 | 0.0811 | 2.3327 | 0.0197 | * | 8 / 4 |
| Pir/Epir | 0.1115 | 0.1327 | 0.8403 | 0.4008 | n.s. | 5 / 4 |
| M1 | -0.0300 | 0.0274 | -1.0962 | 0.2730 | n.s. | 15 / 7 |
| Linear mixed-effects: *ITI prediction* (%) ~ *Task* + (1\|*Session*); reference = Task 1. |  |  |  |  |  |  |

OFC and mPFC supported higher decoding than M1 or Pir/EPir across both tasks, as quantified by the decoder’s log-likelihood (Figure 4B) and prediction accuracy (Figure 4C). Crucially, in the RNNs (Figure 4D-E) and in neural activity from the OFC and mPFC in Task 2 (Figure 4G-H), the decoder increasingly misclassified ISI time points as ITI as the delay progressed, mirroring the declining p(ISI) belief, whereas ITI misclassification remained stable throughout the ISI in Task 1. This divergence was more prominent in the RNNs and in the OFC and mPFC starting ∼2.0 s after cue onset, coinciding with the time at which ISI belief begins to decline in Task 2 (Figure 4F-I), resulting in a significant interaction effect of task and micro-state (Figure 4H). No such difference was found in M1 or Pir/EPir. These effects remained consistent when accounting for the disparities in sample size across brain regions in both cues (Figure S3D). Because decoders were trained on rewarded trials only, with Task 2 omission trials held out for testing, the elevated ITI misclassification in Task 2 cannot reflect a training bias. These results confirm that frontal cortical activity encodes graded belief states rather than ground-truth task states in a task-dependent manner.

Given that, across all levels of analysis, OFC activity consistently matched the predictions of belief-like dynamics, we focused subsequent experiments and in-depth dynamical analyses on this brain region.

### Dynamical system modeling supports the presence of a task-specific ISI attractor in OFC

We next sought to directly characterize the dynamical structure of the population activity in the OFC. To this end, we fit a recurrent switching linear dynamical system (rSLDS)^26,27^ to neural activity recorded in the OFC and in the M1 as a control region, using task events as external inputs *u* (Methods). The rSLDS approximates nonlinear dynamics as a set of switching discrete states *z*_*i*_, each governed by linear dynamics with a single fixed point (Figure 5A). An rSLDS outperformed a standard linear dynamical system (LDS) in capturing OFC activity (Figure S4C), justifying the nonlinear approach. We further evaluated the predictive performance of the rSLDS fits by comparing its forward prediction accuracy with two distinct null models designed to isolate the contribution of across-cell correlations and input-driven dynamics on the model’s prediction accuracy (Methods, Figure S4 E-G). These comparisons are summarized by the scores *ζ*(*tshift*) and *ζ*(*dyn*), respectively, for the example sessions (Methods, Figure S4G). Finally, we assessed the robustness of the inferred dynamics to model initialization by quantifying the reliability of the flow fields across independent initializations (Methods, Figure S4D).

**Figure 5.**
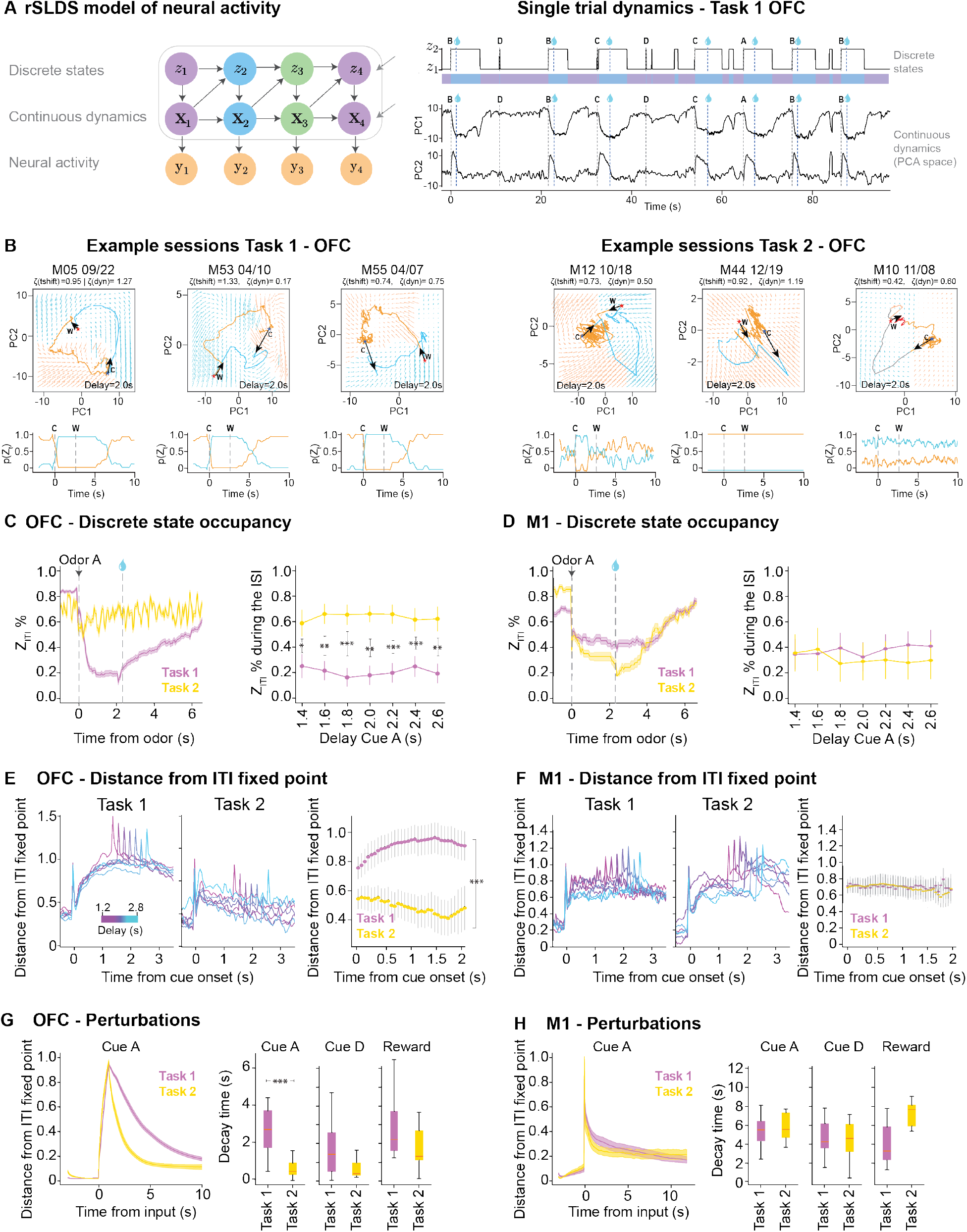
Dynamics of neural activity in the OFC are consistent with dynamics of beliefs and value-RNNs. A. Left: The rSLDS consists of a set of discrete states (*Z*_*i*_), each defining a linear dynamical system (*x*_*i*_) and with a different emission function from the latent dynamics to neural activity (*y*). Right top: rSLDS discrete state occupancy during single trials for an example session in Task 1. Letters A-D indicate the onset of cues presentation; colors indicate occupancy of one of two discrete states. Right bottom: Projection of the rSLDS latent dynamics to the first two PCs for the same single trials. B. Examples of rSLDS fits on single sessions from Task 1 (left) and Task 2 (right). Top: Projection of the rSLDS latent dynamics during Cue A trials (Delay 2s) to the space spanned by the first two PCs. Bottom: rSLDS discrete state occupancy across trials. **C**: cue onset; **W**: water onset. C. Occupancy of *Z*_*ITI*_ from rSLDS fits to neural data in the OFC. Left: Percent *Z*_*ITI*_ occupancy across trials, averaged across sessions for Task 1 and Task 2. Right: *Z*_*ITI*_ occupancy as a function of reward delivery delay (n=21/10 sessions for Task 1/Task 2). Effect of task type computed with a linear-mixed effects model for each delay; dependent: *Z*_*ITI*_occupancy, predictor: task type, random intercept: session. See Table 8 for statistical values. D. Same as panel C but for neural data in M1 (n=11/6 sessions for Task 1/Task 2) E. Left and middle: Distance of rSLDS dynamics from the ITI fixed point in the OFC during the trial session in Task 1 and Task 2. Right: Distance from the ITI fixed point during the ISI period in each task. Interaction effect of task type and time in the ISI delay computed with a linear-mixed effects, dependent: distance from ITI fixed point, predictors: task type and time in the delay, random intercept: session. F. Same as panel E but for M1. G. Left: Distance of rSLDS dynamics from the ITI fixed point following a perturbation input (Cue A, left) in the OFC. Right: Average decay time for the rSLDS latent dynamics to return to the ITI fixed point after input perturbations (Cue A, Cue D, and reward). Effect of task type computed with a linear-mixed effects model for each trial type; dependent: decay time, predictor: task type, random intercept: session. (n=21/10 sessions for Task 1/Task 2). See Table 9 for statistical values. H. Sames a panel G but for M1 (n=11/6 sessions for Task 1/Task 2). Results are represented as mean ± SEM. ***p < 0.001; **p ≤ 0.01; *p ≤ 0.05

**Table 8.** Effect of task on the percentage of occupancy of rSLDS in *Z*_*ITI*_ during the ISI.

|  |  |  |  |  |  |  |
| --- | --- | --- | --- | --- | --- | --- |
| **Delay** | **Coef.** | **SE** | **z** | **p** | **Sign.** | **N /task** |
| 1.4 | 0.4452 | 0.1664 | 2.6748 | 0.0075 | ** | 19/8 |
| 1.6 | 0.3755 | 0.1585 | 2.3693 | 0.0178 | * | 19/8 |
| 1.8 | 0.4128 | 0.1542 | 2.6766 | 0.0074 | ** | 19/8 |
| 2 | 0.3725 | 0.1557 | 2.3915 | 0.0168 | * | 19/8 |
| 2.2 | 0.3424 | 0.1554 | 2.2037 | 0.0275 | * | 19/8 |
| 2.4 | 0.3549 | 0.1660 | 2.1375 | 0.0326 | * | 19/8 |
| 2.6 | 0.2234 | 0.1810 | 1.2344 | 0.2170 | n.s. | 19/8 |
| Linear mixed-effects: $Z_{ITI}$ occupancy (%) ~ Task + (1\|Session); reference = Task 1. | | | | | | |

**Table 9.** Effect of task on the decay time of rSLDS dynamics in response to inputs.

|  |  |  |  |  |  |  |
| --- | --- | --- | --- | --- | --- | --- |
| **Input** | **Coef.** | **SE** | **z** | **p** | **Sign.** | **N /task** |
| Cue A | -3146.415 | 977.978 | -3.217 | 0.001 | *** | 19 / 8 |
| Cue D | -2276.513 | 1222.282 | -1.863 | 0.063 | n.s. | 25 / 8 |
| Reward | -2177.665 | 1367.499 | -1.592 | 0.111 | n.s. | 25 / 8 |
| Linear mixed-effects: $Z_{ITI}$ occupancy (%) ~ Task + (1\|Session); reference = Task 1. | | | | | | |

Our key prediction was that, if OFC encodes belief states, neural activity in Task 1 should be attracted to a separate ISI fixed point distinct from the ITI, and thus, require the system to transition to a distinct rSLDS discrete state, a structure absent in Task 2.

The rSLDS fits of single sessions during Task 1 and 2 showed signatures of distinct dynamical structures (Figure 5B), with a tendency for the latent dynamics during Task 1 to become attracted to two distinct fixed points during the ISI and ITI, but with a tendency to converge to the same fixed point during Task 2 (Figure 5B). To systematically quantify this, we identified the discrete rSLDS state most frequently occupied during the ITI (*z*_*ITI*_) and tracked how often the activity remained in *z*_*ITI*_ over time on a trial-by-trial basis. In Task 1, *z*_*ITI*_ occupancy decreased after cue onset and remained low throughout the ISI, while in Task 2 it initially decreased but gradually recovered as the delay progressed (Figure 5C). This difference in the *z*_*ITI*_ occupancy between tasks was present in OFC but absent in M1. Quantifying the distance of latent trajectories from the ITI fixed point confirmed that the ensemble activity moved progressively further from the ITI fixed point throughout the delay in Task 1, but decayed back toward it in Task 2, in OFC but not in M1 (Figure 5E-F).

Finally, we tested whether the rSLDS dynamics predicted task-specific relaxation times following reward omissions, a signature observed in Value-RNNs but not in echo-state networks^12^. Simulating the forward dynamics after an odor input, Task 1 trajectories took significantly longer to return to the ITI fixed point than Task 2 trajectories in OFC. This difference was not observed in M1, nor for the unrewarded Cue D or reward input (Figure 5G-H). Together, these results support the idea that OFC dynamics exhibit a task-specific ISI fixed point that prolongs activity following cue onset in Task 1, closely mirroring the fixed-point structure predicted by belief-state dynamics and learned by Value-RNNs.

### Omission probe trials confirm task-specific attractor dynamics in OFC

To test the fixed-point predictions empirically without relying on rSLDS fits, we next used reward omission trials as natural perturbations. A separate cohort of mice (n = 4 mice) was first trained on Task 1 and then exposed to unexpected omissions in a "switch session" before transitioning to Task 2 (Figure 6A-B). We compared persistence of neural responses during omission trials in these switch-session mice to omission responses in mice trained exclusively on Task 2.

**Figure 6.**
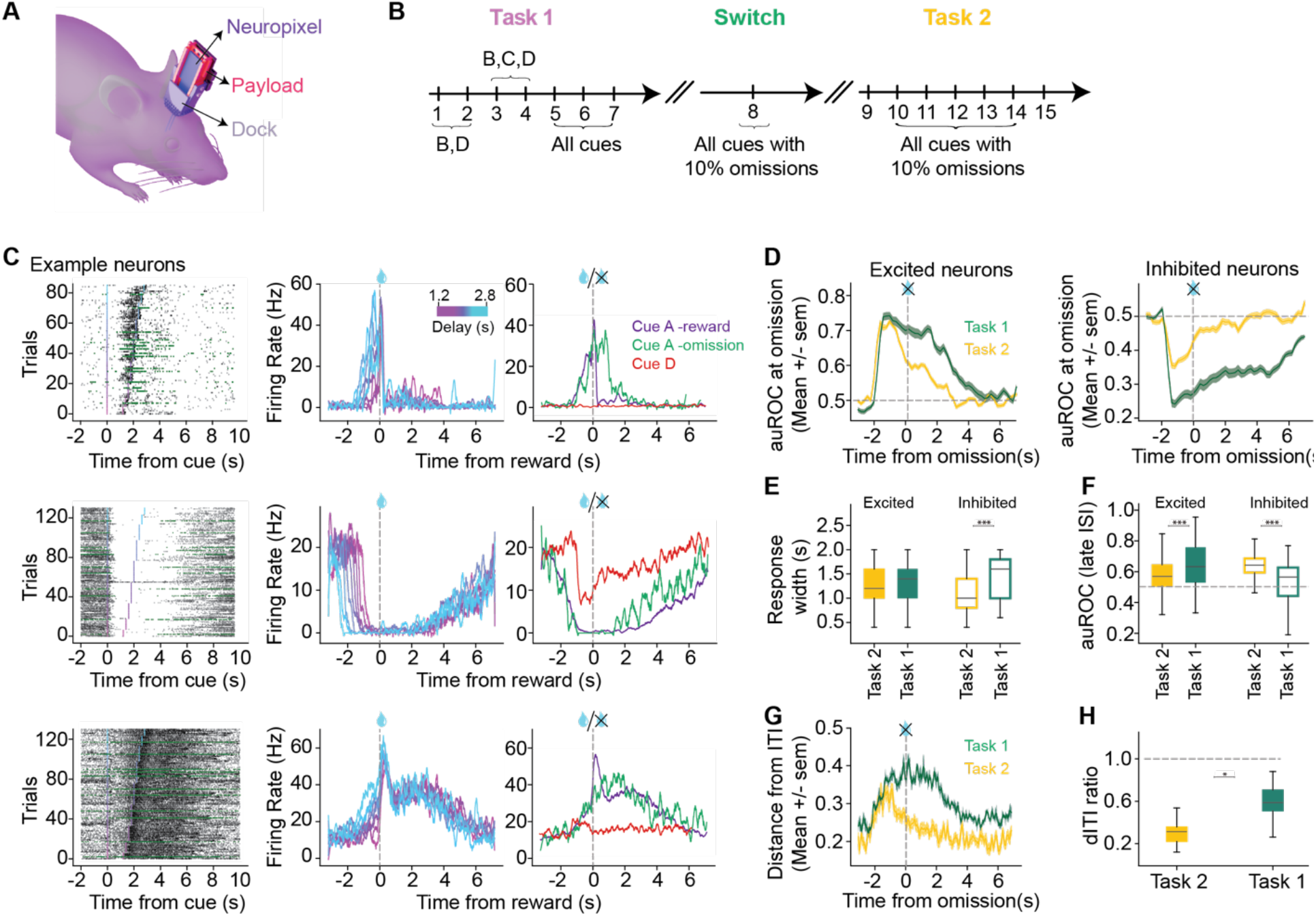
Trajectories of neural activity in response to cues followed by reward omissions follow belief state predictions. A. Schematic of the chronic Neuropixels implants used for the recordings during learning consisting of a ‘payload’ that holds the probe, and a ‘dock’ that keeps the probe stable on the skull. B. Schematic of the training paradigm for the switch experiment. Animals were first trained in Task 1 with the gradual introduction of cues. After 7-9 sessions in Task 1, they underwent the switch session, where reward omissions were introduced. They were then trained on Task 2. C. Spiking activity from three example neurons during the switch session. Left: Raster plots showing spike times across Cue A trials that were rewarded (black) or unrewarded (green). Middle: Average firing rate across all Cue A trials, aligned to the time of reward delivery and plotted as a function of reward delay. Right: Average firing rate across rewarded Cue A trials, unrewarded Cue A trials, and Cue D trials. D. Left: auROC values for neurons that were excited by the cue during omission trials in Cue A, comparing animals trained on Task 1 during the switch session (chronically implanted animals) and those trained on Task 2 (acute recordings) (N=81/457 neurons for Task 1/ Task 2). Right: Same as left panel, but for neurons that were inhibited by the cue (N=54/235 neurons for Task 1/ Task 2). E. Response width distribution based on the auROC from Cue A trials separating excited and inhibited neurons. Same data as in panel D. Effect of Task was computed with linear mixed effects model and pairwise post-hoc comparisons using the Tukey Honestly Significant Difference (Tukey HSD) test. See Table 10 and 11 for exact statistical values. F. Distribution of the auROC during the late ISI taken from Cue A trials separating excited and inhibited neurons. Same data as in panel E. Effect of Task was computed with linear mixed effects model and pairwise post-hoc comparisons using the Tukey HSD test. See Table 12 and 13 for exact statistical values. G. Distance of population activity from the average ITI activity during omission trials in Cue A, for animals trained on Task 1 during the switch session and those trained on Task 2 (N=33 omission trials for Task 1, 75 omission trials for Task 2). H. Percent dITI during omission trials in Cue A. Same data as in panel G. Effect of task type computed with a linear-mixed effects model; dependent: dITI ratio, predictor: task type, random intercept: session (p=0.0214, coefficient for Task 2= -0.2977, SE=0.1294, z = -2.3007) ***p < 0.001; **p ≤ 0.01; *p ≤ 0.05.

**Table 10.**
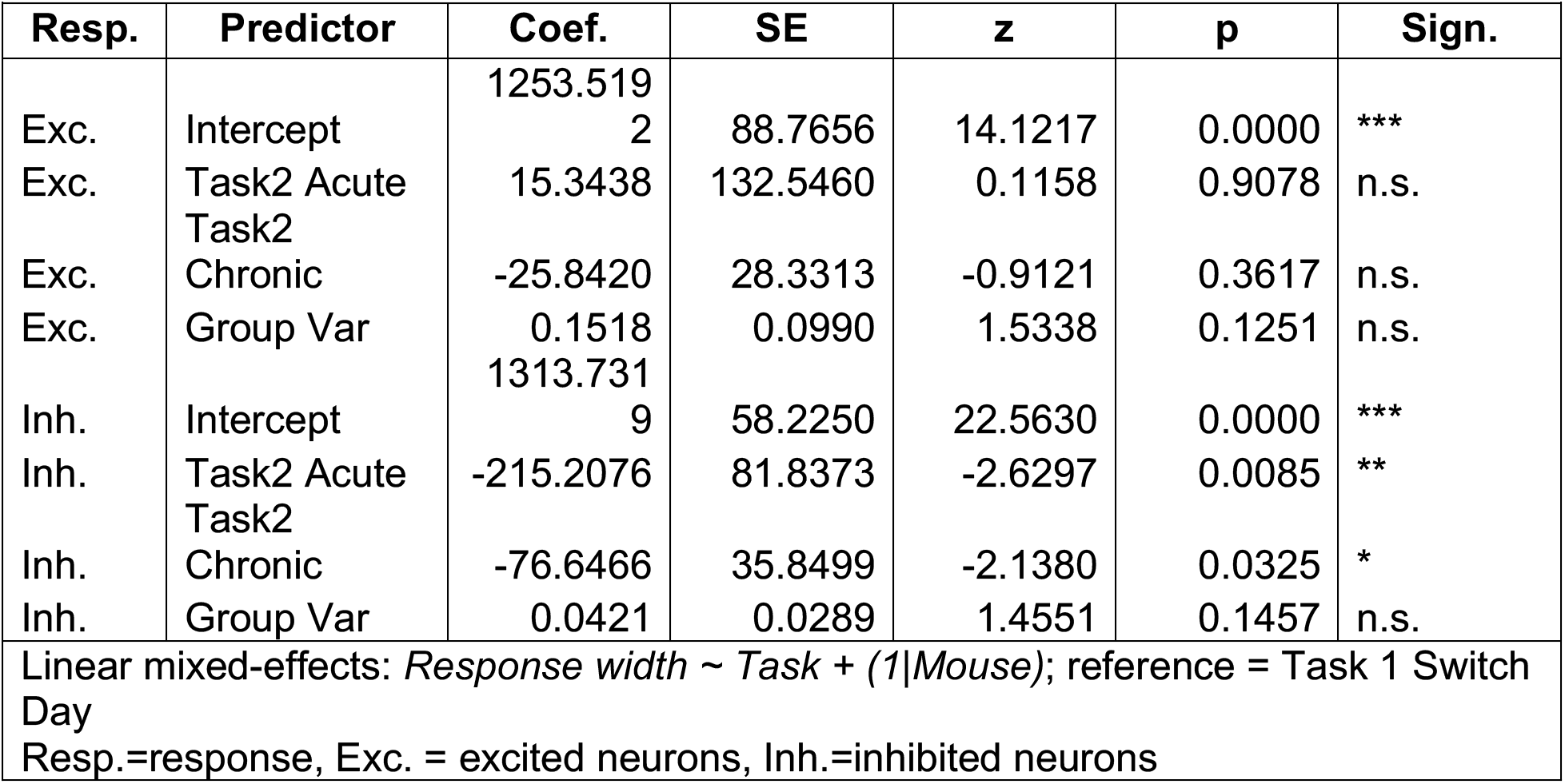
Effect of task on response width of single neurons in omission trials of Cue A.

**Table 11.** Post-hoc pairwise comparison with Tukey HSD for results in Table 10.

|  |  |  |  |  |  |  |  |
| --- | --- | --- | --- | --- | --- | --- | --- |
| **Resp.** | **Group 1** | **Group 2** | **Mean Diff.** | **p-value** | **Lower CI** | **Upper CI** | **Sign.** |
| Exc | Task 1<br>Switch Day | Task2<br>Acute | -63.5327 | 0.0585 | -128.82 | 1.75 | n.s. |
| Inh | Task 1<br>Switch Day | Task2<br>Acute | -276.998 | 0 | -362.47 | -192 | *** |

**Table 12.** Effect of task on auROC at late ISI of single neurons in Cue A omission trials.

| Resp. | Predictor | Coef. | SE | z | p | Sign. |
| --- | --- | --- | --- | --- | --- | --- |
| Exc. | Intercept | 0.6150 | 0.0177 | 34.8082 | 0.0000 | *** |
| Exc. | Task2 Acute | -0.0390 | 0.0257 | -1.5142 | 0.1300 | n.s. |
| Exc. | Task2 Chronic | -0.0203 | 0.0081 | -2.5091 | 0.0121 | * |
| Exc. | Group Var | 0.0656 | 0.0437 | 1.5007 | 0.1334 | n.s. |
| Inh. | Intercept | 0.3649 | 0.0191 | 19.1418 | 0.0000 | *** |
| Inh. | Task2 Acute | 0.0901 | 0.0273 | 3.2986 | 0.0010 | *** |
| Inh. | Task2 Chronic | 0.0227 | 0.0105 | 2.1689 | 0.0301 | * |
| Inh. | Group Var | 0.0586 | 0.0397 | 1.4752 | 0.1402 | n.s. |
| Linear mixed-effects: *auROC at late ISI ~ Task + (1\|Mouse)*; reference = Task 1 Switch Day<br>Resp.=response, Exc. = excited neurons, Inh.=inhibited neurons |  |  |  |  |  |  |

**Table 13.** Post-hoc pairwise comparison with Tukey HSD for results in Table 12.

| Resp. | Group 1 | Group 2 | Mean Diff. | p | Lower CI | Upper CI | Sign. |
| --- | --- | --- | --- | --- | --- | --- | --- |
| Exc. | Task 1<br>Switch Day | Task2<br>Acute | -0.0526 | 0 | -0.071 | -0.03 | *** |
| Inh. | Task 1<br>Switch Day | Task2<br>Acute | 0.1155 | 0 | 0.0904 | 0.141 | *** |

We chronically implanted Neuropixels and recorded OFC activity^29,30^. In switch-session mice, omission trials elicited single-neuron responses that remained elevated well beyond the expected reward time, reflected in significantly higher response widths relative to Task 2-trained mice (Figure 6C-E). Population activity also showed prolonged deviation from the ITI baseline (greater distance from ITI state) extending past expected reward delivery (Figure 6F). Similar results were observed for Cues B and C (Figure S5). These results directly confirm that OFC dynamics learned under Task 1 contingencies support an ISI attractor that sustains neural activity following reward omissions, a signature of task-specific belief-state encoding that is absent in animals trained under Task 2.

### Belief-state dynamics in OFC are absent in naïve animals and emerge through learning

To establish whether belief-state dynamics are acquired through learning rather than present from the outset, we analyzed OFC activity during early training in mice used in the reward omission experiment described above (n = 4 mice)^29,30^. Cues were introduced sequentially: Cues B and D first, then Cue C, then Cue A. Anticipatory licking developed rapidly within sessions (∼20 trials) and generalized across cues, with lick rates for newly introduced Cues A and C exceeding those seen at the equivalent stage of Cue B training (Figure 7A-B).

**Figure 7.**
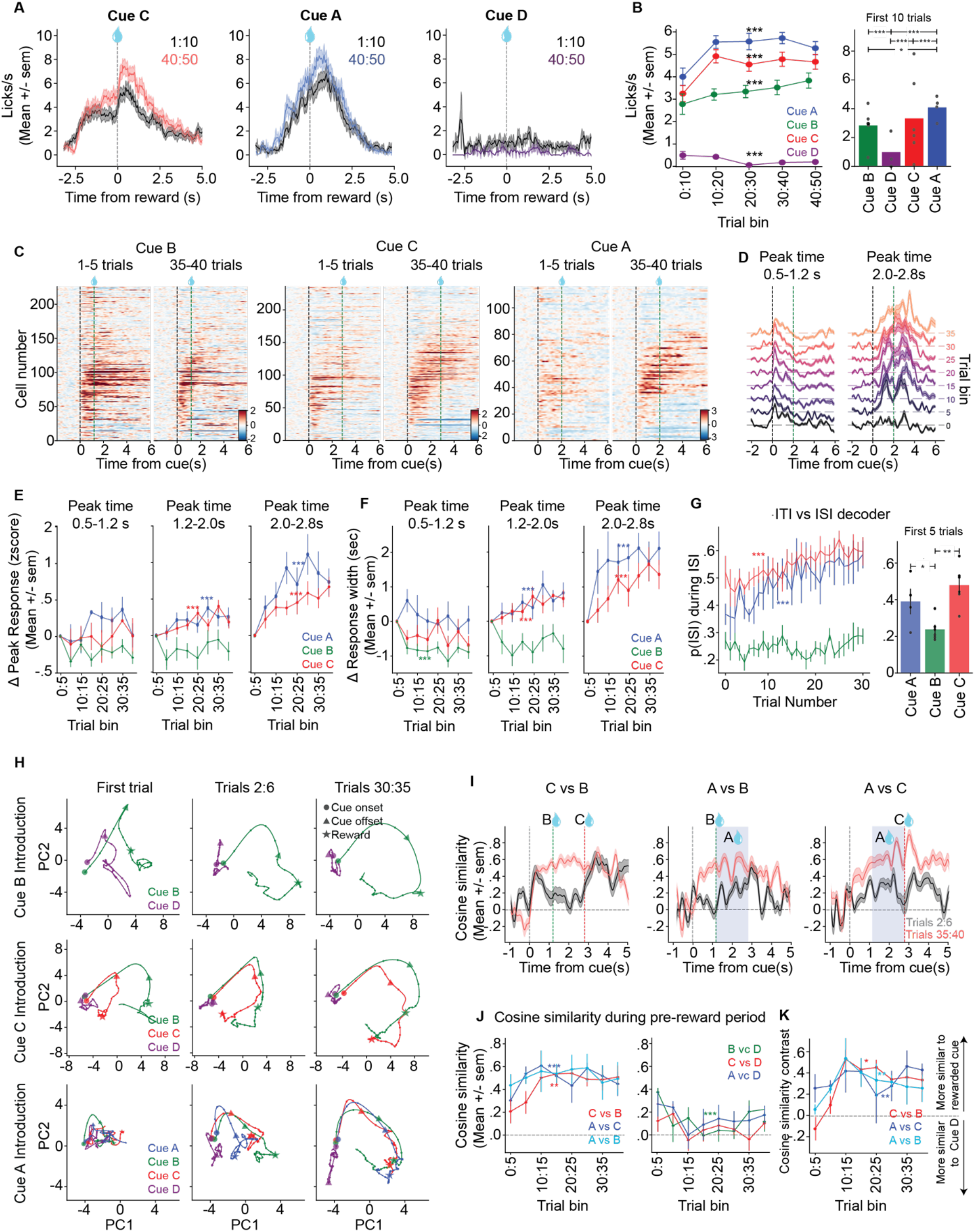
Signatures of belief state encoding in neural activity emerge with learning of the task. A. Anticipatory lick rates for each cue during early (trials 1–10, gray) and late (trials 40–50, colored) training periods. Each panel shows data for one cue. B. Left: Anticipatory licking evoked by each cue as a function of trial number following cue introduction (trials grouped into bins of 10). A linear mixed-effects model (LME) tested the effect of trial bin (continuous predictor) with mouse as a random effect. Right: Mean anticipatory licking during the first 10 trials following cue introduction. To compare lick rates during the pre-reward interval across all pairs of trial types pairwise LME were fit with mouse as a random effect. The full set of pairwise p-values was corrected for multiple comparisons (fdr_tsbky; statsmodels multipletests). See Table 14 for additional statistical details. C. Heatmaps of single-neuron activity evoked by rewarded trial types (Cues B, C, A) during the first five trials after cue introduction (left) and during late training (trials 35–40; right). Neurons were pooled across mice and sorted by the time of peak activity during trials 35–40. Color scale shows z-scored activity. D. Average activity across training (5-trial bins) for neurons responsive during the early ISI (0.5–1.2 s after cue onset; left) and late ISI (2.0–2.8 s; right). E. Change in peak response amplitude (z-score) across training bins for neurons responsive during early (0.5–1.2 s; left), mid (1.2–2.0 s; middle), and late (2.0–2.8 s; right) ISI periods. Effect of trial bin computed with a linear-mixed effects model for each category of neurons and for each trial type; dependent: peak response, predictor: trial bin, random intercept: mouse. See Table 15 for additional details on the statistical values. F. Change in response width across training bins for neurons responsive during early (0.5–1.2 s; left), mid (1.2–2.0 s; middle), and late (2.0–2.8 s; right) ISI periods. Effect of trial bin computed with a linear-mixed effects model for each category of neurons and for each trial type; dependent: response width, predictor: trial bin, random intercept: mouse. See Table 16 for additional details on the statistical values. G. Decoding of task state from population activity. Left: Probability of being in the interstimulus interval (p(ISI)) predicted by a linear decoder as a function of training trial during the first session for each cue. Right: Mean p(ISI) during the first five trials after cue introduction. Effect of trial bin computed with a linear-mixed effects model for each trial type; dependent: p(ISI), predictor: trial bin, random intercept: mouse. To compare p(ISI) across all pairs of trial types pairwise LME were fit with mouse as a random effect. The full set of pairwise p-values was corrected for multiple comparisons. See Table 17 for additional details on the statistical values. H. Population activity projected onto the first two principal components for an example session. Trajectories are shown for the first trial of exposure to each rewarded cue (left), trials 2–6 (middle), and trials 35–40 (right). I. Cosine similarity between population trajectories evoked by each pair of rewarded cues during early training (trials 2–6; gray) and late training (trials 35–40; red). J. Mean cosine similarity during the pre-reward period as a function of training bin between pairs of rewarded cues (left) and between each rewarded cue and Cue D (right). To test whether the metric changes over learning, and given the nonlinearity, a likelihood ratio test (LRT) comparing a cubic spline model against an intercept-only null model was done, p-values are reported in the figure. An additional LRT was done comparing a cubic spline model to a linear model. See Table 18 for additional details on the statistical values. K. Cosine similarity contrast computed as (Cos.Sim (rewarded pair)-Cos.Sim (rewarded-Cue D)/Cos.Sim (rewarded pair)+Cos.Sim (rewarded-Cue D)) as a function of training bin. Positive values indicate greater similarity between rewarded cue trajectories, whereas negative values indicate greater similarity between a rewarded cue and Cue D. To test whether the metric changes over learning, and given the nonlinearity, a likelihood ratio test (LRT) comparing a cubic spline model against and intercept-only null model was done, p-values are reported in the figure. See Table 19 for additional details on the statistical values. ***p < 0.001; **p ≤ 0.01; *p ≤ 0.05.

**Table 14.** Effect of trial of learning on anticipatory licking rate.

| Cue | Coef | SE | z | p | Sign. | N |
| --- | --- | --- | --- | --- | --- | --- |
| A | 0.0278 | 0.0108 | 2.5779 | 0.0099 | ** | 200 |
| B | 0.0237 | 0.0101 | 2.3468 | 0.0189 | * | 298 |
| C | 0.0270 | 0.0091 | 2.9710 | 0.0030 | ** | 300 |
| D | -0.0198 | 0.0074 | -2.6840 | 0.0073 | ** | 143 |
| Linear mixed-effects: *Lick rate ~ Task + (1\|Mouse)* |  |  |  |  |  |  |

**Table 15.**
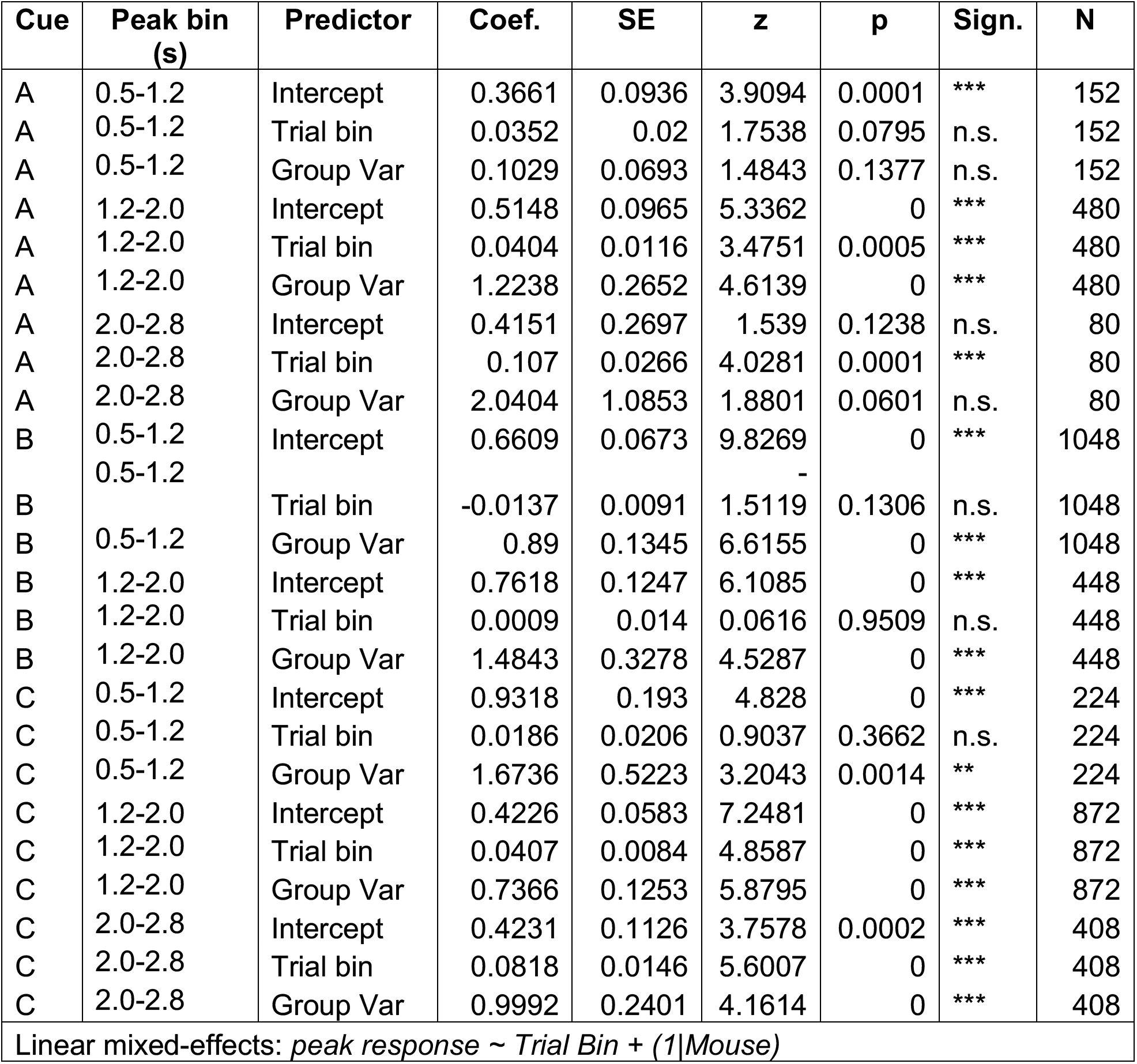
Effect of trial bin of learning on the peak response of single neurons (excited neurons)

**Table 16.**
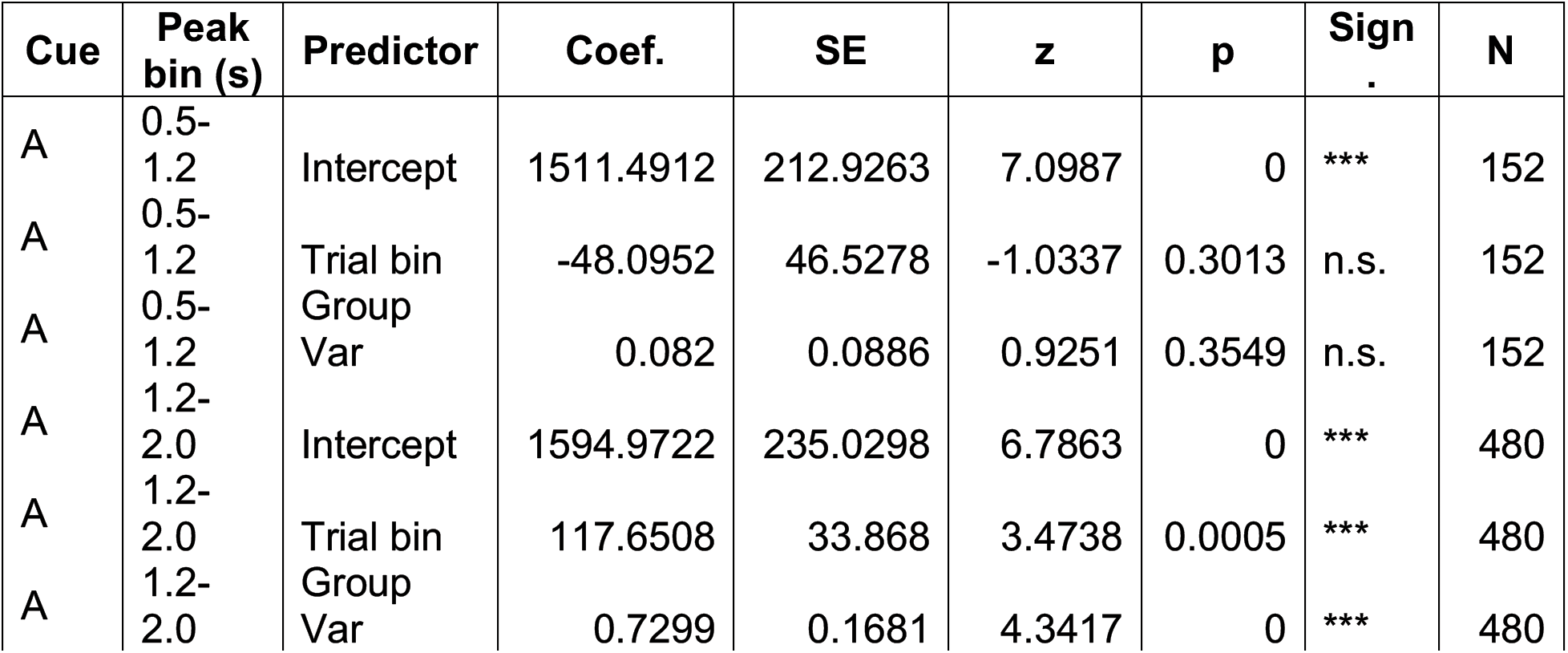
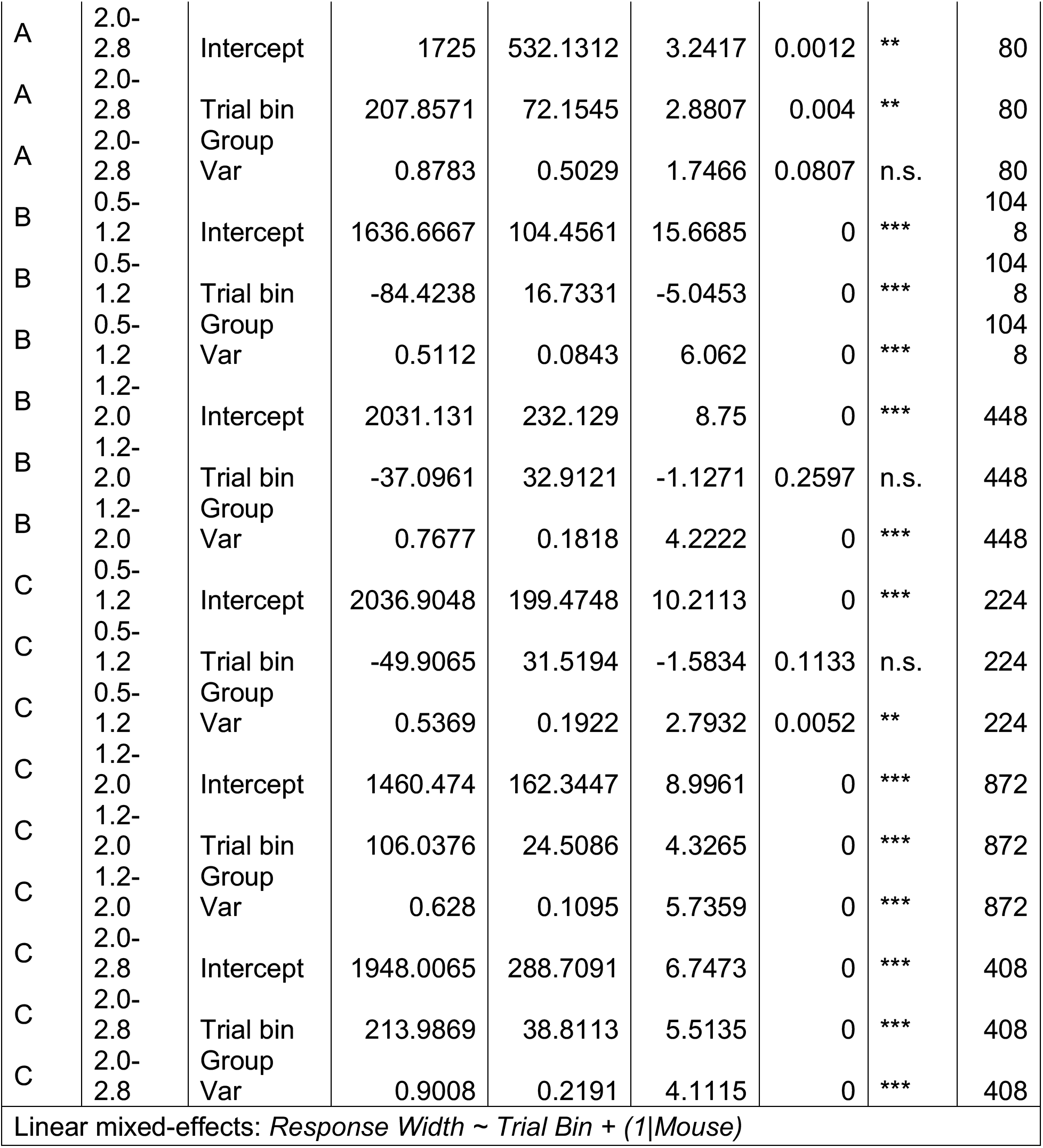
Effect of trial bin of learning on the response width of single neurons (excited neurons)

**Table 17.** Effect of trial of learning on the p(ISI) predicted by a decoder trained in population activity.

|  |  |  |  |  |  |  |
| --- | --- | --- | --- | --- | --- | --- |
| **Cue** | **Coef.** | **SE** | **z** | **p** | **Sign.** | **N** |
| A | 0.0030 | 0.0007 | 4.2369 | 0.0000 | *** | 90 |
| B | 0.0013 | 0.0008 | 1.5214 | 0.1282 | n.s. | 180 |
| C | 0.0043 | 0.0007 | 6.1261 | 0.0000 | *** | 90 |
| Linear mixed-effects: *p(ISI)* ~ *Trial number* + (1\|*Mouse*) |  |  |  |  |  |  |

**Table 18.**
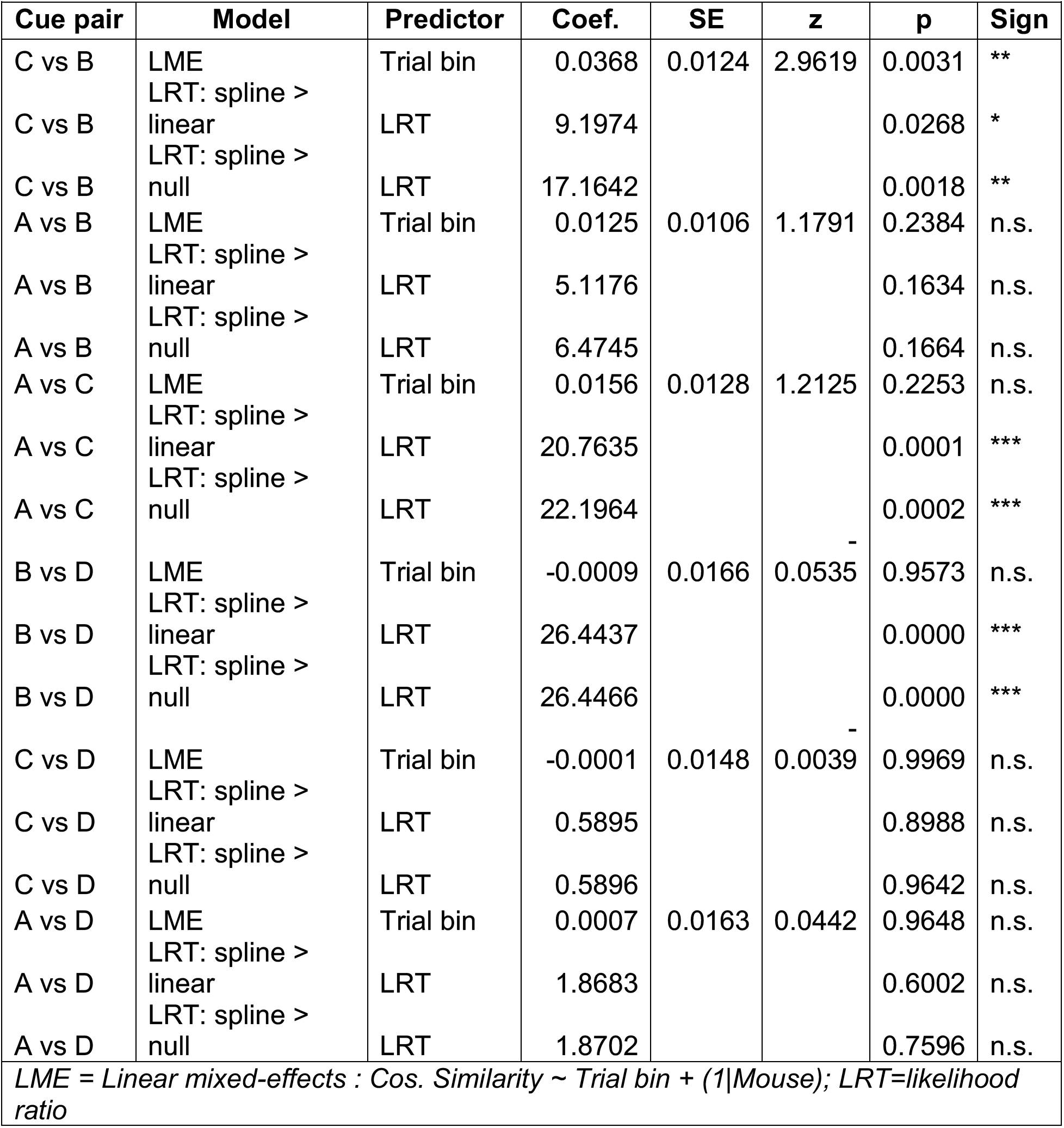
Effect of trial bin on cosine similarity of neural activity between cue pairs.

**Table 19.** Effect of trial bin on cosine similarity contrast of neural activity.

|  |  |  |  |  |  |  |  |
| --- | --- | --- | --- | --- | --- | --- | --- |
| **Cue pair** | **Model** | **Predictor** | **Coef.** | **SE** | **z** | **p** | **Sign.** |
| C vs B | LME | Trial bin | 0.0267 | 0.0162 | 1.6553 | 0.0979 | n.s. |
| C vs B | LRT: spline > linear | LRT | 8.8311 |  |  | 0.0316 | * |
| C vs B | LRT: spline > null | LRT | 11.4854 |  |  | 0.0216 | * |
| A vs B | LME | Trial bin | 0.0266 | 0.0198 | 1.3412 | 0.1799 | n.s. |
| A vs B | LRT: spline > linear | LRT | 12.1043 |  |  | 0.0070 | ** |
| A vs B | LRT: spline > null | LRT | 13.8477 |  |  | 0.0078 | ** |
| A vs C | LME | Trial bin | 0.0323 | 0.0222 | 1.4541 | 0.1459 | n.s. |
| A vs C | LRT: spline > linear | LRT | 11.9352 |  |  | 0.0076 | ** |
| A vs C | LRT: spline > null | LRT | 13.9736 |  |  | 0.0074 | ** |
| LME = Linear mixed-effects : Cos. Similarity Contrast ~ Trial bin + (1\|Mouse) ;<br>LRT=likelihood ratio |  |  |  |  |  |  |  |

At the single-neuron level, early responses to Cue B were broad across the ISI, gradually sharpening into the temporally structured pattern seen after learning (Figure 7C; Figure S6B). For Cues A and C, responses were initially sparse and concentrated near cue onset; with learning, neurons responsive during the mid and late ISI developed progressively larger amplitudes and wider response widths (Figure 7C-F; Figure S6). These learning-related changes were concentrated in late-ISI neurons, and were absent in early-ISI neurons and in responses to Cue B, consistent with the cue-specific development of sustained activity.

At the population level, ISI vs. ITI decoding accuracy improved with learning for Cues A and C, and generalization was evident: decoders assigned higher ISI probability to early Cue A and C trials than to early Cue B trials (Figure 7G). PCA trajectories showed that rewarded cue responses became progressively more aligned with one another during the late ISI while diverging from Cue D, as quantified by cosine similarity (Figure 7H-J). The time of peak activity was strongly correlated across rewarded cues but not with Cue D (Figure S6G–H).

Together, these findings show that belief-state encoding in OFC is absent in naïve animals and emerges through learning, ruling out explanations based solely on sensory-driven responses or intrinsic OFC circuit dynamics. Moreover, the progressive alignment of population trajectories across rewarded cues suggests that learning promotes generalization across task states.

## Discussion

In this study, we examined the neural mechanisms underlying the representation and updating of belief states. We recorded ensemble neural activity from OFC, mPFC, and other regions in mice performing two variants of a Pavlovian task that differ only in reward reliability, producing distinct within-trial belief dynamics. Our results show that ensemble activity in OFC, and to a lesser extent mPFC, exhibits attractor-like population dynamics that closely mirror the task-specific belief dynamics predicted by normative models and by Value-RNNs trained via TD learning. Consistent with belief-state encoding, a linear decoder trained on OFC and mPFC activity could classify task states with accuracy that tracked the animal’s moment-to-moment belief uncertainty: in the partially observable task, ITI misclassification during the ISI increased progressively over the delay, resembling the decay of ISI belief, whereas decoding remained stable in the fully observable task. These neural signatures were absent in motor and olfactory areas, could not be attributed to overt behavioral differences between tasks, and emerged progressively with learning. Furthermore, the slow, attractor-like dynamics were confirmed in the reward omission experiment in mice pre-trained in the deterministic task (Task 1). Together, these findings provide a biologically plausible neural implementation of time-evolving belief-state inference, supporting the dynamical systems framework for implementing belief states, proposed in prior modeling work^8,12^.

A central concept in reinforcement learning is the state, the internal representation of environmental conditions, on which value and policy are computed. Despite its importance, the nature of state representations remains poorly understood. In classic TD learning models, the brain is assumed to maintain a set of temporal "microstates" or basis functions spanning the delay between a cue and reward^3,31–33^. This framework has successfully accounted for many features of dopaminergic responses, including the negative dip upon reward omission and the transfer of responses from reward to cue during learning^31,34^. However, these models rest on biologically untenable assumptions: the state space must be fully specified before any learning takes place, every sensory cue must trigger its own reproducible set of microstates spanning all possible delays, and these representations must be stable across repeated exposures^35,36^. Furthermore, previous studies of dopamine assumed hand-crafted state representations to explain non-canonical dopamine responses as TD errors^3–6,8,15,16,32,37,38^. However, how these states are acquired and implemented in neural circuits remains unknown.

Our work takes steps toward demystifying these unknowns. Rather than assuming a pre-specified state space, we show that the brain can learn task-dependent state representations through experience: OFC develops task-specific attractor dynamics that naturally encode the statistics of within-trial state transitions, reflecting the degree of uncertainty due to partial observability. Traditional TD models address timing by postulating fixed temporal basis functions—microstates or microstimuli—that tile the cue-reward interval and must be assigned to individual cues before learning^32,33^. Our dynamical systems framework avoids these limitations. Rather than pre-specifying a state sequence, the brain can use recurrent dynamics that naturally approximate the task structure and the progression of time within a specific task (also see, ^36^).

A fundamental challenge in reinforcement learning is that sensory input typically provides only partial information about the current state of the world: the true underlying states are often hidden and must be inferred. A principled solution is to replace deterministic (but partially observed) state representations with belief states: probability distributions over possible states that are continuously updated as new observations arrive^1,2,11^. Even in simple Pavlovian tasks, stochasticity in task events such as probabilistic reward omissions introduces partial observability, necessitating belief-state inference^3,4^. In the present study, we exploited this structure, contrasting fully deterministic and partially observable reward schedules, to produce tasks with qualitatively distinct belief dynamics^4^. Our results show that these distinct dynamics are reflected in OFC population activity as attractors that differ in their “leakiness”: in the deterministic task, a stable ISI attractor sustains ensemble activity away from the ITI baseline throughout the delay, whereas in the probabilistic task, the absence of reward causes activity to drift gradually back toward the ITI state, reflecting the declining belief of being in the ISI. Under the dynamical systems framework, the belief state – the probability distribution over ITI and ISI – is therefore encoded not in the activity of individual neurons, but in the location of the population activity along the trajectory between the two task-state attractors.

In summary, the present study applies the dynamical systems view to explain how the brain represents task structures and belief states. Our results indicate that the brain solves the problem by learning a dynamical system whose attractor structure reflects the statistical regularities of the environment, a solution that is both flexible and scalable in ways that classical models are not.

## Acknowledgements

We thank David M. Zoltowski for advice on the modelling of neural data using rSLDS; Takahiro Yamaguchi for initial work in modelling the task with RNNs; and Soichiro Doi for contributing with pilot analysis. We also thank the instructors from the Cajal School for Machine Learning in Neuroscience, including Laura Driscoll, Memming Park and Alex Cayco-Cajic for advice on the scope of using dynamical systems for understanding neural computations. We thank all the Uchida lab members for helpful comments at the different stages of this work.

## Author contributions

### Data and code availability

Data and code will be posted to online repositories upon publication

## Methods

### EXPERIMENTAL MODEL AND SUBJECT DETAILS

#### Mice

We used 16 adult male mice ranging in age from 6 to 12 months of age (C57/BL6). Animals ranged in weight from 20-25 g. Animals were singly housed on a 12-h dark/12-h light cycle (dark from 7AM to 7PM). We trained animals on the behavioral task at approximately the same time each day, between noon and 7PM. The recordings were made at the same times as their training. All experiments were performed in accordance with the National Institutes of Health Guide for the Care and Use of Laboratory Animals and approved by the Harvard Institutional Animal Care and Use Committee.

### METHOD DETAILS

#### Surgical procedures

Before training, mice were implanted with a headplate and ground pin. We performed all surgeries under aseptic conditions while mice were anesthetized with isoflurane (inducted with 4%, maintained at 1.5% in oxygen, 1 L/min). Buprenorphine (SR 0.6 mg/kg, intraperitoneal injection) was administered to provide analgesia. The mice were then moved to a stereotaxic frame. Local anesthetic (bupivacaine) was applied to the scalp to cover the incision site and the eyes were lubricated with ointment (Puralube). Body temperature was maintained at 37°C using a feedback-controlled heat pad (Harvard Apparatus). A custom headplate and ground pin were implanted as previously described. Briefly, the stereotaxic frame was adjusted to ensure bregma and lambda were in the same horizontal level. The skull surface was etched for 30 seconds (Enamel Etchant Gel; C&B-Metabond) before the headplate was affixed by dental cement (C&B -Metabond). A small craniotomy was drilled in the posterior skull, which was used for the insertion of a ground pin that was also affixed by dental cement.

For mice undergoing acute recordings, we used a permanent marker to mark the target coordinates on the skull along the prefrontal cortical regions (AP: +1.0 to +2.5, ML: 0.5. to 2.0) For mice undergoing chronic recordings, a craniotomy of approximately 0.6 mm in diameter was drilled at the target site (OFC: AP +2.5, ML 1.0). The craniotomy and the skull were then covered with silicone sealant (KwikCast, World Precision Instruments). Mice were left to recover inside their home cages on top of a heating pad.

#### Behavioral training

After 1 week of post-surgical recovery, mice were water-restricted in their cages. Weight was maintained above 85% of pre-restriction body weight. We habituated and briefly head-restrained mice for 2-3 days before training. Odors and task events were delivered to animals with using custom-made hardware (Bpod, Sanworks). Each odor was dissolved in mineral oil at 1/10 dilution. 30mL of diluted odor was placed into glass fiber filter-paper, and then diluted with filtered air 1:20 to produce a total 1L/min flow rate.

Mice were trained in a Pavlovian associative learning task used in previous studies (Starkweather et al. 2017; Starkweather, Gershman, and Uchida 2018). All cues were odors that were presented for 1 second. Cue A predicted reward delivery at a variable delay sampled from a discrete Gaussian distribution (t_A_ = {1.2,1.4,1.6,1.8,2.0,2.2,2.4,2.6,2.8}s from cue onset), Cues B and C at a short and long delay t_B_ =1.2, t_C_=2.8 from cue onset), respectively, and Cue D predicted no reward. In Task 1, Cues A-C were always followed by a reward, whereas in Task 2, reward was omitted in 10% of trials.

The odors used for training were: s(-) limonelle, 1-fenchone, 1-4 cineole, 1-hexanol, octanoic acid, ethyl-acetate and eugenol. The odor-cue assignment was randomized across mice.

The training paradigm consisted of progressively adding Cues based on difficulty. First Cue B and D, then Cue C and finally Cue A. The total training time lasted from 2 to 3 weeks.

#### Acute Neuropixels electrophysiological recordings

After mice had achieved good task performance, we performed acute electrophysiological recordings in their prefrontal cortical regions. Craniotomies were performed the day before recording and covered with Kwik-Cast (World Precision Instruments), and mice were allowed to recover overnight.

Recordings were made using Neuropixels probes (Imec) and spiking data was acquired custom-written software (http://billkarsh.github.io/SpikeGLX/). Neuropixels probes were lowered using a Thorlabs micromanipulator (PT1-Z8) at 9 μm/sec. After reaching the target depth, the probe was allowed to settle for 20-30 minutes prior to starting the recording. Recordings lasted an hour on average and were terminated if the mouse was no longer engaging with the task with a maximum number of 300 trials in each session.

#### Chronic Neuropixels electrophysiological recordings

We printed the parts of the Neuropixels implants in the Harvard Neurotech core (https://www.ntcore.org/) using a resin based SLA (stereolithography) 3D printer (Formlabs). The parts were based on a previously published design (Bimbard et al. 2024) but adapted so that it was suitable to be implanted in the OFC. The Neuropixels probe was mounted on the custom-made parts: a ‘payload’ supports the probe while a ‘dock’ protects the probe shaft and it is the part that is attached to the skull.

The probe implantation was performed at least five days post the initial surgery. Mice were anesthetized with ketamine/dexmedetomidine (60 mg/kg, intraperitoneal injection) and headfixed using the headplate previously implanted. The silicon sealant was removed, and the area cleaned with saline. The probe was inserted slowly using a micromanipulator (Luigs & Neuman) into the craniotomy until reaching the desired depth (4 mm DV). Artificial dura (Cambridge Neurotech) was applied on the craniotomy around the probe shank. The dock of the implant was then cemented to the skull with two layers of cement, waiting 15 minutes between layers to allow for setting. Once the second layer dried, the probe holder was detached from the probe implant and the flex cable was secured on the implant using a ZIF connector (FH26W-45S-0.3SHW, Hirose Electric Co) attached to the payload. After implantation, the anesthesia was then reversed using atipamezole (Antisedan, 0.5 % mg/kg, intraperitoneal injection).

#### Spike sorting

Neuropixels data was spike sorted offline with Kilosort 3, followed by manual curation in Phy (https://github.com/kwikteam/phy). During manual curation, each cluster of spikes detected in Kilosort was compared to those with similar waveform shapes to determine whether they should be merged. The merging step was done considering: spike waveform similarity, drift patterns, and cross-correlogram features.

### QUANTIFICATION AND STATISTICAL ANALYSIS

#### Decoding analysis for task type based on lick rates

To elucidate whether the temporal structure of the licking patterns differed between the two tasks we trained a logistic regressor using licking rates (100 ms time bins) as regressors and task type as the variable to predict (Python function ‘sklearn.linear-model.LogisticRegression’). We compared the prediction accuracy against the one of a null model where the labels had been shuffled.

#### Area under the receiver operating characteristic curve (auROC)

For computing the auROC for each single neuron we followed what was done by (Cohen JY, Haesler S, Vong L, Lowell BB, Uchida N. 2012) and compared the histogram of spike counts during the baseline period (pre-cue period) to that during a given bin within the trial by moving a criterion from zero to the maximum firing rate for a given neuron. We then plotted the probability that the activity during a given time bin was greater than the criteria against the probability that the baseline activity was greater than the criteria. The area under this curve quantifies the degree of overlap between the two spike count distributions. An auROC value close to 0.5 implies activity close to baseline, auROC>0.5 indicates increased activity with respect to baseline, and auROC <0.5 decreased activity with respect to baseline.

Neurons were classified as excited or inhibited based on the predominant direction of their discriminative responses during the ISI. For each neuron, we applied two thresholds to the AUROC timecourse: excitatory threshold (auROC ≥ 0.6, indicating elevated firing), inhibitory threshold (auROC ≤ 0.4, indicating suppressed firing in the target condition). For each threshold, we binarized the auROC values during the ISI period (+1 if exceeding threshold, -1 otherwise) and computed the mode (most frequent value) across time bins. Neurons were classified as: ‘excited’ if the mode of the binarized auROC at the 0.6 threshold was positive and ‘inhibited’, if the mode of the binarized auROC at the 0.4 threshold was negative.

For each neuron we then computed the response width as the number of time bins during the ISI where auROC >0.6 or <0.4 for excited and inhibited neurons, respectively. The response width represents the cumulative duration (in milliseconds) that a neuron maintains discriminative activity above threshold during the ISI. Higher response widths indicate neurons with sustained selective responses, while lower values indicate transient activity.

##### Statistical tests

To compute the effect of the task type on each brain region on the auROC during the late ISI or the rauROC, we performed a linear-mixed effects model having as predictor the task type (Task 1 or 2) and the mouse as a random intercept to account for inter-subject variability (Python function ‘statsmodels.formula.api.mixedlm’)

#### Projection of neural activity to the ISI/ITI -dividing plane with Linear Discriminant Analysis

Neural data (20 ms time bins) was organized into a 3D array with dimensions (neurons x trials x time). Firing rates for each neuron were z-scored across both time and trials relative to a pre-cue baseline period (-2 to 0 s relative to cue onset).

For each trial type we constructed training data to discriminate between the ISI (from cue to reward delivery) and ITI periods (from -1.5s before cue onset to cue onset). For rewarded trial types, only trials where reward was delivered were included in the training set. Neural activity was reshaped into a 2D matrix where each time point from each trial became a separate sample, resulting in dimensions (samples × neurons). Binary labels were assigned to each sample (0 = ISI, 1 = ITI). We concatenated data from all trial types to allow the classifier to learn a common ISI/ITI discriminant axis that generalized across different odor cues and delay conditions.

We then fit a Linear Discriminant Analysis model using singular value decomposition using (sklearn.discriminant_analysis.LinearDiscriminantAnalysis with solver=’svd’). For binary classification, LDA identifies a single discriminant axis that maximally separates the two classes by maximizing the ratio of between-class to within-class variance.

To visualize neural trajectories along the learned discriminant axis, we projected all neural onto this axis. The projection value represents the signed distance of sample *I* along the axis that maximally separates ISI from ITI states. Higher (more positive) values indicate the sample is more "ITI-like," while lower (more negative) values indicate it is more "ISI-like."

#### Principal component analysis (PCA)

We performed principal component analysis (PCA) on trial-averaged population activity. For each neuron, spike counts were first averaged across trials within each condition (trial type or Cue A delay) and then concatenated, yielding a matrix of Nneurons x Nconditions x T (where T is the number of time bins per condition). Each neuron’s activity was then z-scored across the concatenated condition–time axis. PCA was performed treating each time point as an observation and neurons as features. Low-dimensional trajectories were obtained by projecting the z-scored activity onto the leading principal components. We quantified the variance explained by each component using the eigenvalue spectrum and computed cumulative explained variance (Figure S1C). Population dimensionality was further summarized using the participation ratio: 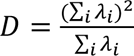 where *λ*_*i*_ is the ith PCA eigenvalue. This measure was additionally normalized by the number of recorded neurons to facilitate comparisons across sessions.

#### Fraction of explained variance by trial average responses

For each neuron, we computed the trial-averaged activity across trials at each time point, yielding a mean temporal response profile. The fraction of explained variance (FVE) for each neuron was computed as:

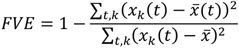

Where *x*_*k*_(*t*) denotes the activity on trial k at time t, *x̅*(*t*) is the trial averaged activity at time t, and *x̅* is the overall mean across trials and time. The denominator is the total variance: the sum of squared deviations of single-trial activity from the neuron’s overall mean (computed across all trials and time points). The residual variance is the numerator: the sum of squared deviations of single-trial activity from the trial-averaged response at each time point. This measure captures the extent to which trial-to-trial variability is accounted for by the condition-locked mean response, with higher values indicating more reliable, stimulus-locked activity.

#### Decoding analysis for task states

For the state decoding analysis, we followed^12^. The goal here is to test whether the potentially belief-like representations in neural activity are sufficient to decode the underlying state. Thus, we trained a linear decoder to infer the underlying true state *s*_*t*_ ∈ {1, …, *K*} (defined above) using a linear transformation of the neural activity.

We first z-scored the activity of each neuron to have zero mean and unit variance. We then performed a multinomial logistic regression. After training, the decoder’s estimated state probabilities over *s*_*t*_ are:

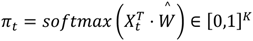

Where:

- *Ŵ* ∈ ℝ^*N*+1×*K*^ contains the decoder parameters
- *X*_*t*_ ∈ ℝ^*N*+1^ is the neural activity at time *t* after standardization, plus an extra constant column of 1’s to fit the offset
- Softmax function normalizes the vector to be a valid probability over the *K* values of *s*_*t*_

To evaluate the resulting decoder, we calculated the model’s log-likelihood (*l*) on the test set as follows:

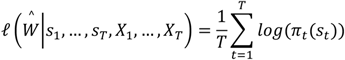

where *π*_*t*_(*s*_*t*_) ∈ [0,1]is the *s*^*th*^ entry of the vector *π*_*t*_.

For Task 2, we left out the omission trials from the training set to avoid artificially biasing the ITI prediction of the decoder trained in Task 2 with respect to the one trained in Task 1.

##### Statistical tests

To test for the effect of task type in the accuracy or rate of ITI prediction for each micro-state we performed a permutation test (1000 iterations) for the difference of means of variable of interest (accuracy or rate of ITI prediction) between the tasks. We used the function ‘scipy.stats.permutation-test’ with 1000 permutations and the mean of the distributions as the test statistic, To test for the effect of task type or brain region in the log-likelihood of the decoder we implemented a linear-mixed effects model with mouse as a random intercept.

#### Distance from ITI analysis

As a linear approach to quantify the persistence in activity at a population level without relying on any dimensionality reduction method, we computed the vector norm distance (Euclidean distance) between the population activity at a given time bin (*X*_*t*_) and the mean position of the population activity during the pre-cue ITI period (*X*_*ITI*_) defined as:

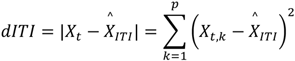

The level of persistence in activity was computed by taking the ‘dITI ratio’ defined as:

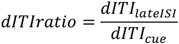

Where:

- *dITI*_*lateISI*_ is the mean dITI during the late ISI period, defined as the time range from 200 ms before reward delivery to reward delivery time.
- *dITI*_*cue*_ is the peak dITI during the cue response, defined as the time range from 0 to 500 ms from cue onset.

##### Statistical tests

To test for the effect of task type in the distance from the ITI during the late ISI period and dITI ratio metrics we implemented a linear-mixed effects model with mouse identity as a random intercept.

#### Recurrent switching linear dynamical systems (rSLDS)

To infer potential mechanisms that generated the observed ensemble neural activity, we fitted an rSLDS model to the neural activity using the publicly available software developed by Linderman et al., (https://github.com/lindermanlab/ssm, ^26,39,40^). Briefly, an rSLDS is a generative model that approximates a non-linear time series with a sequence of linear dynamical systems. It contains two sets of latent variables: continuous variables *x*_*t*_ and discrete states *z*_*t*_.

The transitions between discrete states are a function of the previous state, external inputs, and the dynamics:

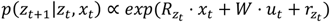

where:

- *R*_*z_t_*_ ∈ ℝ^*K*−1×*D*^ is the weight matrix of recurrent dependencies between continuous states and a discrete state *K*
- *W* ∈ ℝ^*K*−1×*M*^ are the input weights influence on the discrete state transitions,
- *r*_*z*_*t*__ ∈ ℝ^*K*−1^ is the bias that captures Markovian dependency of *z*_*t*−1_ on *z*_*t*_

The continuous dynamics are determined by the discrete state at a given time point and are also influenced by external inputs:

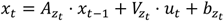

where:

- *A*_*z*_*t*__ ∈ ℝ^*D*×*D*^ is the dynamics matrix for a given discrete state *z*_*t*_
- *V*_*z*_*t*__ ∈ ℝ^*D*×*M*^ are the input weights on the continuous dynamics for a given discrete state *z*_*t*_
- *b*_*z*_*t*__ ∈ ℝ^*D*^ is the bias for a given discrete state *z*_*t*_.

Finally, the model’s emissions are a function of the continuous dynamics with a predefined noise distribution:

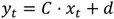

where:

- *C* ∈ ℝ^*N*×*D*^ is a low to high dimensional mapping
- *d*: follows a Poisson distribution in our case.

We fitted rSLDSs to neural activity during task performance at a session-level. Our set of inputs were the four odors and reward, encoded as a one-hot vector with 5 dimensions, and the emissions corresponded to the neural activity recorded from a given brain region. The hyperparameters of the model are the number of discrete states (*K*) and number of continuous dimensions (*D*). These hyperparameters were selected independently for each session using a grid search over K= {2,3,4,5,6} and D = {2,3,4,5,6,7,8,9,10,12,14,16,18}. For each K and D combination, the model was fit and its evidence lower bound (ELBO) was evaluated on the held-out test set. We selected the smallest K-D pair whose test-set ELBO was within 5% of the maximum ELBO obtained across all tested combinations for that session (Figure S4A). The selected hyperparameters were then used for all subsequent analyses of that session (Figure S4B).

#### Model comparison: rSLDS vs LDS fits

To elucidate whether the dynamics of neural activity in the OFC are explained by a linear or a non-linear dynamical system, we fitted a linear dynamical system (LDS) and a nonlinear dynamical system (rSLDS) to all the sessions performed in Task 1 and 2 in the OFC. The LDS was defined as an rSLDS but with the number of discrete states *K* = 1. Both models incorporated task events as external inputs.

To compare model performance, we quantified the accuracy with which the inferred latent dynamics predicted neural activity forward in time on held-out data. For each session, the model dynamics were initialized at −200 ms relative to cue onset, and model emissions were generated forward in time including the task inputs. At each time point, forward predictive accuracy was quantified as the Pearson correlation coefficient between the predicted neural population activity and the observed neural activity. Correlations were computed separately for the pre-cue period (−0.2 to 0 s relative to cue onset) and three post-cue periods (0–0.5 s, 0.5–1.0 s, and 1.0–1.5 s after cue onset). For each session, predictions were evaluated on 500 held-out test trials, and the mean Pearson correlation across trials was used as the session-level measure of forward predictive accuracy (Figure S4C).

#### Goodness of fits: null-models comparison

To determine which components of the rSLDS contributed to its forward predictive performance, we compared the full fitted models with two null models designed to isolate the contributions of static across-neuron correlations and learned dynamics.

##### Circular time-shift null model

This null model controlled for predictive accuracy arising from static correlations across neurons rather than from temporal dynamics. For example, if one neuron is consistently active while another is consistently inactive, the predicted and observed population vectors may exhibit a high correlation at each time point even if the model does not capture the temporal evolution of neural activity. To account for this contribution, for each cue and trial, the observed population activity vector at each time point was correlated with the model-predicted population vector at a circularly shifted time point. Forward predictive accuracy was then calculated using the same Pearson correlation procedure described above.

##### Ablated-dynamics null model

The second null model was designed to assess the contribution of the learned recurrent dynamics to forward predictive accuracy and to determine whether predictive performance could instead be explained primarily by the task driven dynamics. For this model, the dynamics matrix was replaced with a matrix of zeros while all other model parameters were kept unchanged. This procedure removed the contribution of the learned recurrent dynamics while preserving the effects of the external inputs and other components of the fitted model. Forward predictions were then generated and evaluated using the same procedure as for the full rSLDS.

For both null models and the fitted rSLDS, forward predictions were generated from the same initial condition (−200 ms relative to cue onset) and evaluated on held-out data. Session-level predictive accuracy was calculated as the mean Pearson correlation across 500 test trials. Unless otherwise indicated, analyses shown in the figures report the session-level averages for Cue A.

To quantify the deviation of the full model with respect to the nulls we computed the *ζ*(*tshift*) and *ζ*(*dyn*) score defined as:

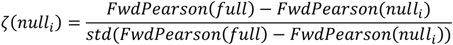

Where:

- *null*_*i*_: corresponds to the time-shifted or dynamics-ablated models for the *ζ*(*tshift*) and *ζ*(*dyn*) scores respectively.
- *full*: corresponds to the full model
- *Fwdlearson*: corresponds to the forward predictive accuracy of each model.

#### Estimation of fixed points of the rSLDS intrinsic dynamics

To estimate the fixed point of the *z*_*ITI*_we solved the dynamics equation for the corresponding discrete state. Given the parameters *A* = *A*_*zITI*_ and *b* = *b*_*zITI*_ and the dynamics equation *x*_*t*+1_ = *A* ⋅ *x*_*t*_ + *b*_*t*_. A fixed point is the point *x*^∗^ such that: *x*_*t*+1_ = *x*_*t*_ = *x*^∗^. Thus:

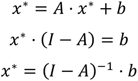

#### Estimation of intrinsic and input-driven dynamics in the rSLDS fit

To simulate the effect of input perturbations in the rSLDS we first initialized the network at fixed point of the discrete state mostly occupied in the ITI (*z*_*ITI*_). The input-driven dynamics are given by:

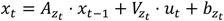

Where *u*_*t*_ is a time-dependent input (5 dimensional for 4 odors and a reward) and *V*_*z*_*t*__ is the weight of the inputs on the dynamics. For performing the perturbations we assigned *u*_*t*,*i*_ = 1 to the *i*_*th*_ input of interest and plotted one update step, giving a ‘snapshot’ into the input-driven dynamics elicited by the input onset.

#### Input perturbations in the rSLDS fits

For performing input perturbations, we took the following steps:

1. We initialized the dynamics at the identified fixed point for the *zITI* (*x∗_z_ITI__*).
2. We designed input perturbation *ut* of each cue or reward. The vector *ut* is a time-dependent input (5 dimensional for 4 odors and a reward). For performing the perturbations, we assigned *ut*,*i* = 1 to the *ith* input of interest for a duration of 100 ms, starting 1 second after initialization.
3. We ran the rSLDS dynamics forward with zero noise for a duration of 20 sec following the dynamics: *xt* = *Az*_*t*_ ⋅ *xt*−1 + *Vz*_*t*_ ⋅ *ut* + *bz*_*t*_ and *p*(*z*_*t*_+1|*z*_*t*_, *xt*) ∝ *exp*(*Rz*_*t*_ ⋅ *xt* + *W* ⋅ *ut* + *rz*_*t*_).
4. To quantify the decay times to the fixed points, we computed the pre-stimulus baseline, and defined the decay time as the time point at which the norm-distance from that baseline crosses the zero distance.

##### Statistical tests

To test for the effect of task type in the decay time to omission probe we implemented a linear-mixed effects model with mouse as a random intercept and task as a predictor for each cue.

#### Analysis during learning: Computation of peak response time, response sign and width

Given that analysis during learning required following activity across small sets of trials we took a distinct approach from the auROC analysis above to estimate peak response times and response widths during learning. Spike density functions were first computed from binned spike counts (20 ms bins) during the first session of learning a given cue type by first averaging spikes across the last 20 trials of the session and then normalizing firing rates relative to a pre-cue baseline period (-2 to 0 s relative to cue onset) to convert firing to a z-score. The resulting z-scored activity was smoothed using a Gaussian kernel (σ = 100 ms).

Peak response times were computed from the SDF within the cue-to-reward interval. The first time bin crossing a z-score threshold of 0.2 was used as the response onset. Within the response period following this onset, the time of the largest absolute response was identified. The sign of the response (excitatory or inhibitory) was defined as the sign of this extremum. The peak response time was then calculated as the center of mass of the response: 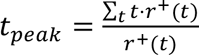 where *r*^+^(*t*) denotes the positive portion of the response after thresholding (values below 0.2 were set to zero).

Response width was quantified around the peak response time using the smoothed z-scored SDF averaged across sets of 5 trials. For each neuron, the SDF was first converted to a unipolar signal by multiplying by the response sign, so that inhibitory responses were treated equivalently to excitatory responses. We defined a threshold of 0.1 z-score units. Time bins exceeding this threshold were identified, and connected components were computed to determine contiguous regions of activity. The region containing the peak response was selected, and the response width was defined as the duration between the first and last time bins in this contiguous region.

#### Analysis during learning: Cosine similarity analysis of population responses

To quantify the similarity between population responses evoked by different cue types, we compared trial-averaged population activity vectors across training using cosine similarity as a metric. Analyses were performed separately between rewarded cues and between rewarded and unrewarded cues.

For each cue pair, we analyzed neural activity during the first session in which the newly introduced cue was presented. Trials were grouped into non-overlapping bins of 5 consecutive trials, spanning trials 0–40 of training. For each trial bin, spike trains were assembled across all recorded neurons. For each neuron and trial type, spike counts were converted to z-scored spike density functions (SDFs) as explained above.

At each time point, the activity of all neurons was represented as a population vector. Cosine similarity between the two trial types was then computed pointwise in time as 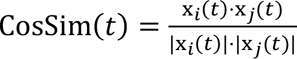 where x*_i_* (*t*) and x*_j_* (*t*) are the population response vectors at time t for trial types i and j, respectively. Thus, cosine similarity quantified the angular alignment between population trajectories independent of their overall magnitude. To obtain stable estimates and account for unequal neuron counts across conditions, we used bootstrap resampling. For each cue pair and trial bin, 2,000 resamples were generated by sampling 100 neurons with replacement from the pooled population. Cosine similarity was computed separately for each resample. This was visualized in Figure 7i.

For analyses comparing alignment among rewarded cues versus alignment between rewarded and unrewarded cues, cosine similarity was averaged within the pre-reward period and then summarized as a contrast index: 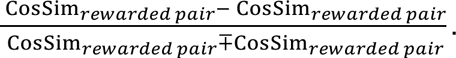

Positive values indicate that trajectories evoked by rewarded cues were more similar to each other than to the unrewarded cue, whereas negative values indicate greater similarity between a rewarded cue and the unrewarded cue.

##### Statistical tests

To test whether similarity and the contrast changed systematically across learning, we quantified the evolution of cosine similarity across trial bins for the average cosine similarity in the pre-reward period (300 ms before reward). We tested for a monotonic change across trial bins using the Spearman rank correlation between trial bin index and the observed similarity values. Cosine similarity was expected to increase across learning for comparisons between rewarded cues, whereas the expected direction was reversed for comparisons involving the unrewarded cue. To assess significance, we generated a null distribution by permuting the order of trial bins independently for each bootstrap resample and recomputing the Spearman correlation. The permutation p-value was defined as the fraction of null correlations greater than or equal to the observed correlation (with direction adjusted according to the predicted learning effect).

#### Analysis during learning: Decoding of task state from population activity

To assess whether OFC population activity improved its representation of task state with learning, we trained a logistic regression classifier on a binary classification (ISI vs ITI). This was performed separately for each cue using pseudopopulations constructed by pooling neurons across sessions and mice.

For training, single-trial spike counts were truncated to a common number of trials across sessions and restricted to the window from -2 to 8 s after cue onset. Three temporal segments were defined for each trial: a pre-cue ITI (-2 to 0 from cue onset), the ISI (cue onset to reward onset) and a post-ISI ITI period (reward onset to 8 s) and each time bin was assigned a label indicating ISI (1) or ITI (0). Each feature corresponded to the zscored activity of each neuron. Decoder performance on the training set was evaluated using 5-fold stratified cross-validation.

To obtain time-resolved state probabilities predicted by the logistic classifier, we used a pseudopopulation resampling procedure. For each cue type, 100 neurons were sampled with replacement from the pooled population, and this was repeated 1,000 times. For each resample, the decoder was trained on late-training trials (the last 20 trials of the session) and tested on the remaining earlier trials producing, a time-resolved estimate of p(ISI) for every trial. To summarize learning-dependent changes, we computed the mean predicted probability of the ISI state during the pre-reward period. This is visualized in Figure 7G.

##### Statistical tests

To test whether state decoding changed across learning, we repeated the same decoding analysis separately for each mouse without pseudopopulation resampling. For each mouse and cue type, the decoder was trained on a late-training window (trials 30–40) and then tested on earlier trials (trials 0–30) and computed the mean predicted probability of the ISI (p(ISI)) during the pre-reward period. We then implemented a linear-mixed effects model with mouse as a random intercept and trial as a predictor of p(ISI). Pairwise comparisons between cue types were then performed to test for differences in p(ISI) between cue types during the first trials of training using two-sided Mann–Whitney U tests corrected for multiple comparisons.

## Supplementary notes

### Belief state inference can be formulated as a dynamical system

Our previous studies ^4^ proposed that our task can be formulated as a semi-Markov environment with two discrete “states”: the inter-stimulus interval (ISI) and the inter-trial interval (ITI). In this formulation, we assume the mouse is estimating a distribution over these two states that varies over time (the “belief state” distribution (*b*_*t*_) Following previous work^4,12,15^, we formalized the process of hidden-state inference by assuming that the ISI and ITI comprise temporal “micro-states” (200 ms time bins, Figure 1A, Methods), and, thus, the belief state distribution has a dimensionality equal to the number of micro-states. Our primary goal is to elucidate how the process of belief state inference is implemented in the brain. An important insight came from our previous study^12^, where it was pointed out that the changing belief states across time can be formulated as a dynamical system. As mentioned earlier, belief states are defined as the posterior probability distribution over states, given the history of observations. This distribution changes over time governed by the following equation:

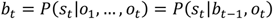

The second equality comes from the Markovian assumption of the state transitions that makes each state transition dependent only on the previous state and allows us to “summarize” the history of observations in the previous belief state *b*_*t*−1_.

Now consider the final conditional probability written in the equation above. This is a probability density function that can be considered as any other function, but that needs to be normalized to sum to 1. Thus, the update equation above could be rewritten more generally as:

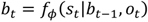

Where now the value of the belief at time *t* is a function, *f*_*ϕ*_(⋅), of its value at the previous time step. Thus, the equation above is the definition of a dynamical system in discrete time, with *f*_*ϕ*_(⋅) the function that determines the evolution of the state variable of the system: the belief state distribution. This dynamical system formulation makes the notion of belief states more generalizable as it doesn’t assume any predefined state structure^2^.

## Supplementary figures

**Figure S1.**
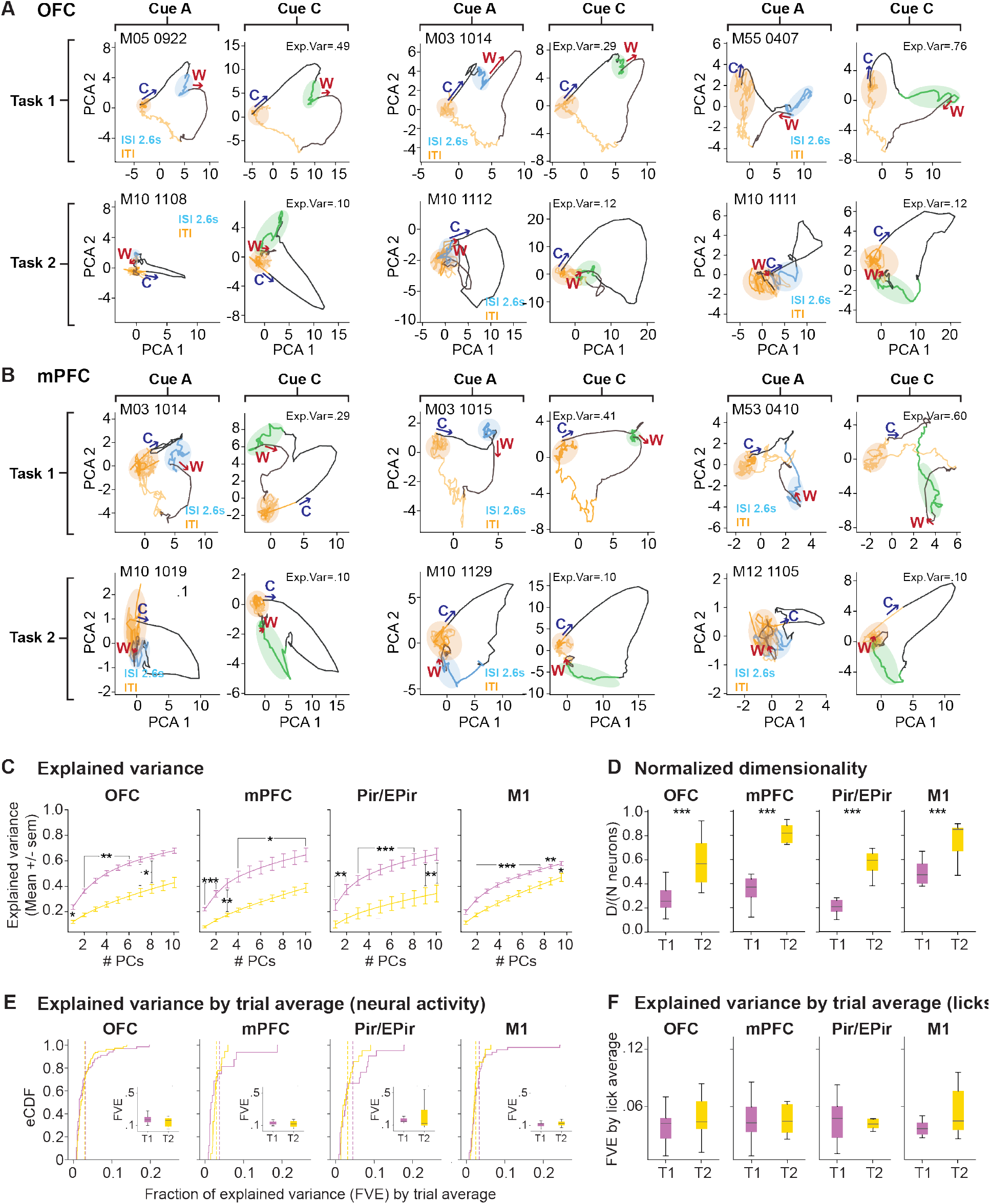
Population activity in frontal cortical areas is consistent with dynamics of beliefs and value-RNNs. A. Trajectories of the first two PCA components in neural activity in the OFC and mPFC for example sessions. Each panel shows Task 1 (top) and Task 2 (bottom) for Cue A at the delay condition of 2.6 s and Cue C. The arrows show the direction of the changes in trajectory elicited by the cue (C) and reward (W). The proportion of variance explained by the first two components is indicated. B. Same as panel A but for neural activity from mPFC. C. Fraction of explained variance of neural activity as a function of the number of PCs considered for Task 1 and 2 in the OFC, mPFC, Pir/EPir, and M1. D. Normalized dimensionality of neural activity in Task 1 and 2 in the OFC, mPFC, Pir/EPir, and M1. Effect of task type computed with a linear-mixed effects model for each delay; dependent: dimensionality, predictor: task type, random intercept: mouse. E. Distributions of fraction of variance of neural activity explained by the trial average in Task 1 and 2 in the OFC, mPFC, Pir/EPir, and M1. Effect of task type computed with a linear-mixed effects model for each delay; dependent: dimensionality, predictor: task type, random intercept: mouse. F. Distributions of fraction of variance of neural activity explained by the trial average of licks in Task 1 and 2 in the OFC, mPFC, Pir/EPir, and M1. Effect of task type computed with a linear-mixed effects model for each delay; dependent: dimensionality, predictor: task type, random intercept: mouse. Results are represented as mean ± SEM. ***p < 0.001; **p ≤ 0.01; *p ≤ 0.05.

**Figure S2.**
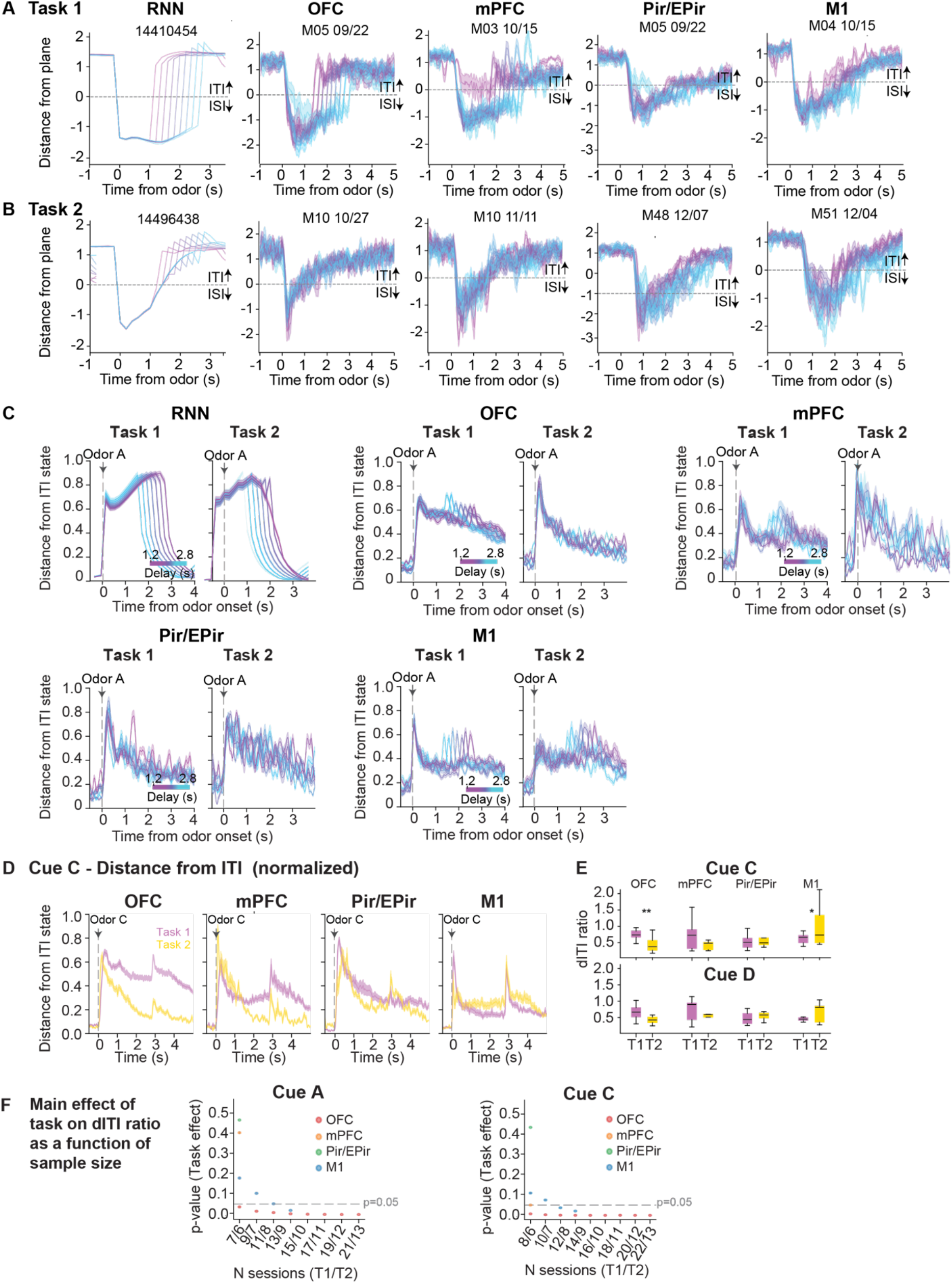
Population activity in frontal cortical areas is consistent with dynamics of beliefs and value-RNNs in rewarded cues. A. Population activity derived from the RNN and neural activity in the OFC, mPFC, Pir/EPir areas and M1 in Task 1 (top) and Task 2 (bottom) was projected to the LDA ISI/ITI separating plane. Traces are shown for example networks and sessions for each brain region averaged across trials for each Cue A delay in Task1. B. Same as panel A but for Task 2. C. Distance from ITI state defined as the Euclidean distance between neural population activity at a given time bin (20 ms) and the mean activity during the ITI, computed for each Cue A delay and averaged across trials in Task 1 and 2. D. Distance from ITI state computed for Cue C in Task 1 and 2. E. dITI ratio derived from neural activity during Task 1 and Task 2 for Cue C and D. Effect of task type computed with a linear-mixed effects model for each delay; dependent: dITI ratio, predictor: task type, random intercept: mouse, corrected for multiple comparisons. F. Mean effect of task on the dITI ratio (p-value from a linear mixed-effects model with session as a random factor) as a function of sample size. At each sample size, an equal number of sessions per brain region were drawn without replacement (n = 10,000 resamples), spanning the range from the smallest to the largest dataset available per brain region. Results are represented as mean ± SEM. ***p < 0.001; **p ≤ 0.01; *p ≤ 0.05.

**Figure S3.**
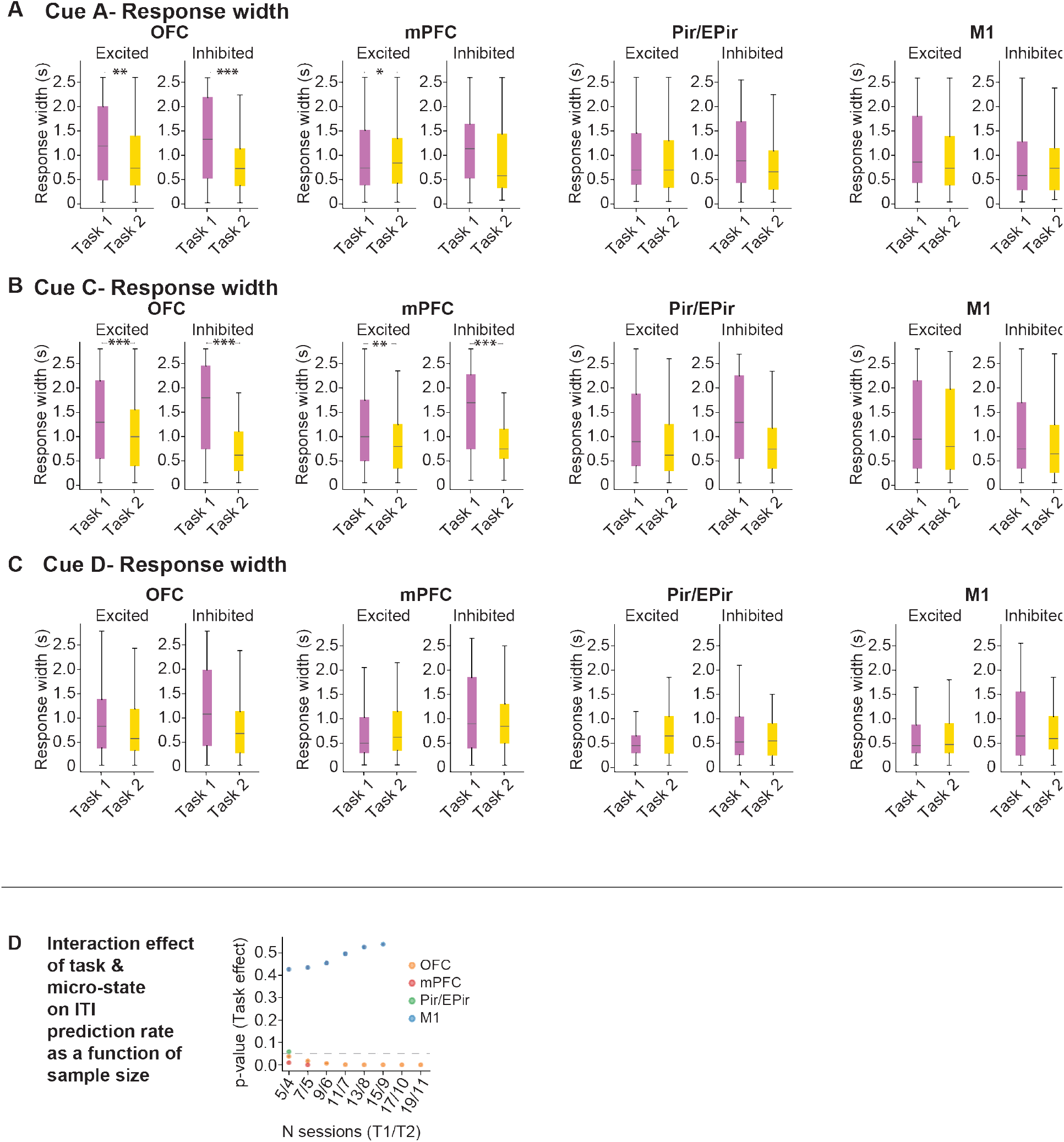
Persistent activity at the population level is supported by prolonged single neuron activity in neurons that are excited or inhibited by the trial period. A. Response width based on the auROC of neural activity from neurons excited and inhibited by the trial period from Cue A in Task 1 and Task 2 in the OFC, mPFC, M1 and Pir/EPir. Effect of task type computed with a linear-mixed effects model; dependent: response width, predictor: task type, random intercept: session. B. Same as panel A but for Cue C. C. Same as panel A but for Cue D. D. Interaction effect of task and micro-state on the ITI prediction rate (as in Figure 4H, p-value from a linear mixed-effects model with session as a random factor) as a function of sample size. At each sample size, an equal number of sessions per brain region were drawn without replacement (n = 10,000 resamples), spanning the range from the smallest to the largest dataset available per brain region. ***p < 0.001; **p ≤ 0.01; *p ≤ 0.05.

**Figure S4.**
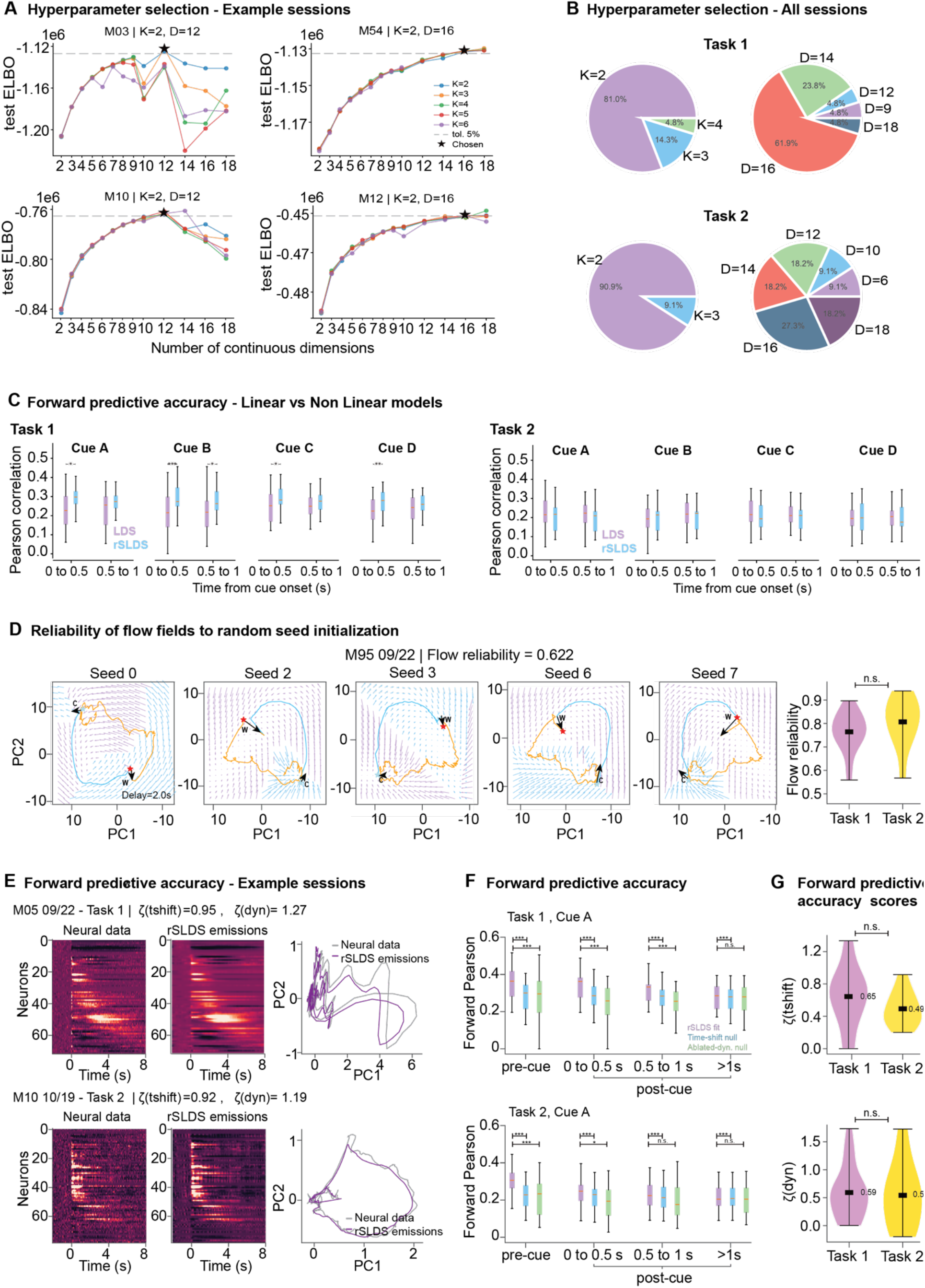
Hyperparameter selection, goodness of fits and model comparison of the rSLDS fits to neural activity. A. Hyperparameter selection for four example sessions. The smallest K–D set with a test-set ELBO within 5% of the maximum ELBO was selected (Methods). B. Distribution of hyperparameters selected across all sessions for Task 1 (top) and Task 2 (bottom). C. Comparison of linear dynamical system (LDS) and recurrent switching linear dynamical system (rSLDS) fits for both tasks. Forward predictive accuracy was quantified as the Pearson correlation coefficient between neural activity and model emissions on unseen test data during the inter-stimulus interval (ISI; 0– 0.5 s and 0.5–1 s from cue onset), following initialization of the dynamics at −200 ms relative to cue onset. The effect of model type was assessed using a linear mixed-effects model with Pearson correlation as the dependent variable, model type as the fixed effect, and session as a random intercept D. Left, example flow fields of the latent dynamics projected onto the space spanned by the first two principal components (PCs) for rSLDS fits obtained with different random seed initializations. Right, distribution of the flow reliability metric across sessions for Task 1 and Task 2. Flow reliability was quantified as the cosine similarity between latent-dynamics flow fields obtained from different seed initializations (Methods). E. Example sessions showing ground-truth neural activity and rSLDS emissions on an unseen test set. Left, single-neuron activity and model emissions, sorted according to the peak time of neural activity relative to cue onset. Right, population activity projected onto the space spanned by the first two PCs. F. Comparison of forward predictive accuracy between the rSLDS and two null models. The time-shift null model accounts for cross-cell static correlations that can increase model accuracy without relying on the intrinsic dynamics of neural activity. The ablated-dynamics null model quantifies the contribution of input-driven dynamics to forward predictive accuracy (Methods). G. Distribution of forward predictive accuracy scores for rSLDS fits to neural activity during Task 1 and Task 2. For each session, the score directly compares the forward predictive accuracy of the full rSLDS model with that of each of the two null models (Methods). ***p < 0.001; **p ≤ 0.01; *p ≤ 0.05.

**Figure S5.**
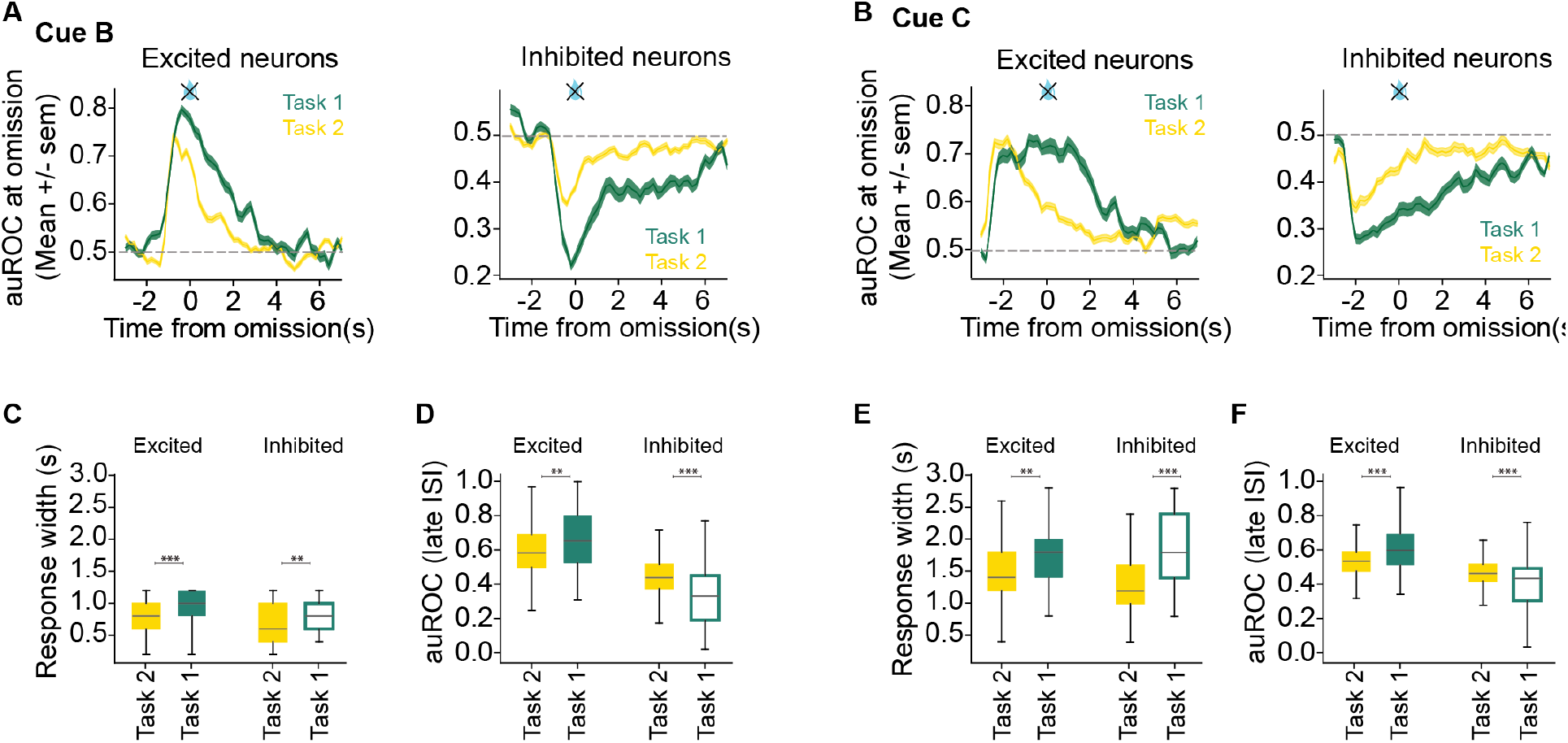
Trajectories of neural activity in response to cues followed by reward omissions follow belief state predictions in rewarded cues. A. Left: auROC values for neurons that were excited by the cue during omission trials in Cue B, comparing animals trained on Task 1 during the switch session (chronically implanted animals) and those trained on Task 2 (acute recordings) (N=81/457 neurons for Task 1/ Task 2). Right: Same as left panel, but for neurons that were inhibited by the cue (N=54/235 neurons for Task 1/ Task 2). B. Same as panel A but for Cue C (N=81/457 neurons for Task 1/ Task 2). C. Response width distribution based on the auROC from Cue B trials separating excited and inhibited neurons. Same data as in panel A. Effect of task type computed with a linear-mixed effects model for each trial type; dependent: response width, predictor: task type, random intercept: session D. Distribution of the auROC during the late ISI taken from Cue B trials separating excited and inhibited neurons. Same data as in panel A. Effect of task type computed with a linear-mixed effects model for each trial type; dependent: auROC during the late ISI, predictor: task type, random intercept: session E. Same as panel C but for Cue C. Same data as in panel B. Effect of task type computed with a linear-mixed effects model for each trial type; dependent: response width, predictor: task type, random intercept: session F. Same as panel D but for Cue C. Same data as in panel B. Effect of task type computed with a linear-mixed effects model for each trial type; dependent: auROC during the late ISI, predictor: task type, random intercept: session Results are represented as mean ± SEM. ***p < 0.001; **p ≤ 0.01; *p ≤ 0.05.

**Figure S6.**
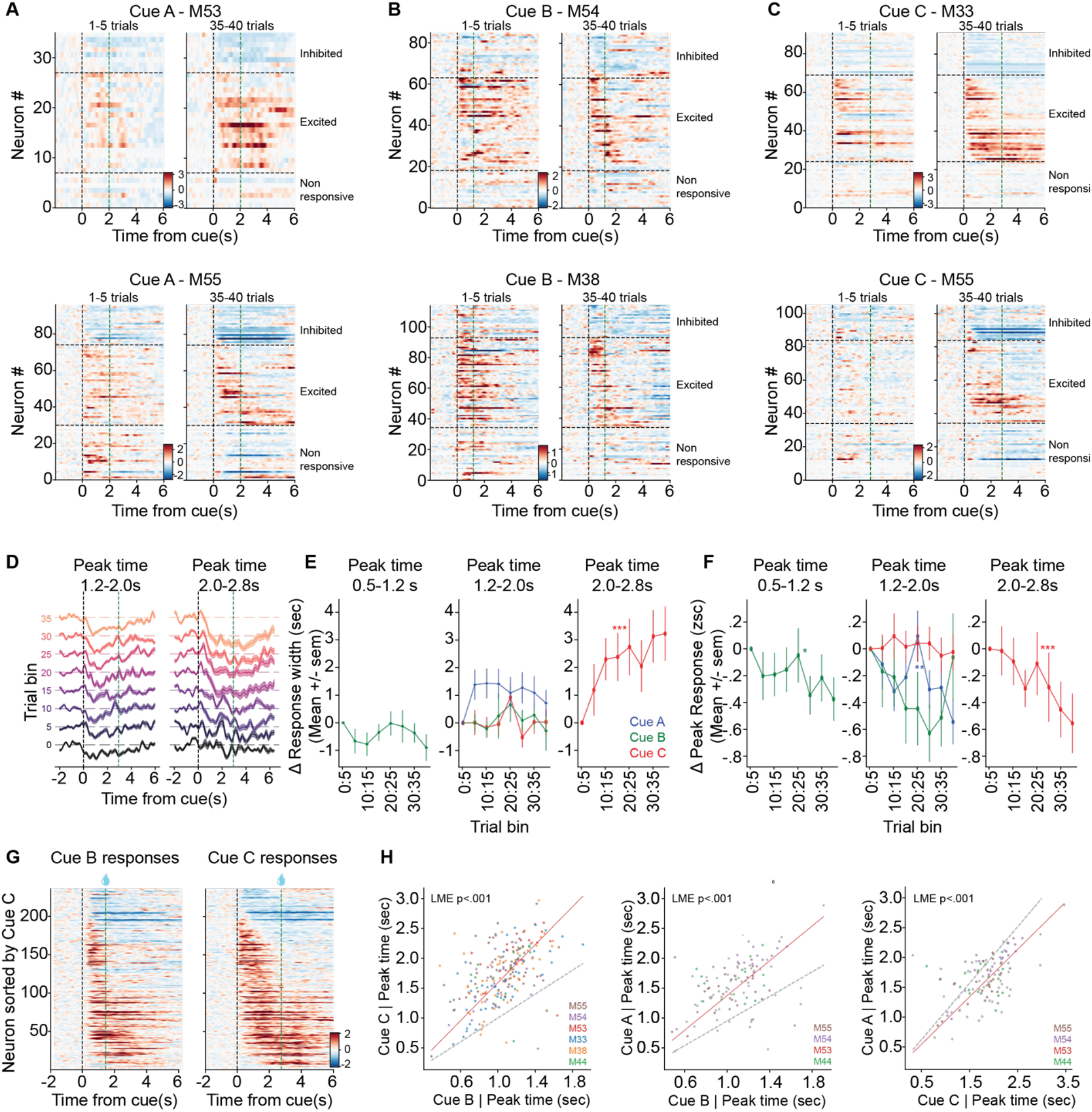
Signatures of belief state encoding in neural activity emerge with learning of the task. A. Heatmaps of single-neuron activity evoked by Cue A during the first five trials after cue introduction (left) and during late training (trials 35–40; right). Neurons sorted by the time of peak activity during trials 35–40 for two example sessions. Color scale shows z-scored activity. B. Same as panel A but for Cue B. C. Same as panel A but for Cue C.D. Average activity across training (5-trial bins) for neurons responsive with inhibition during the mid ISI (1.2-2.0 s after cue onset; left) and late ISI (2.0–2.8 s; right). E. Change in response width across training bins for neurons responsive with inhibition during early (0.5–1.2 s; left), mid (1.2–2.0 s; middle), and late (2.0–2.8 s; right) ISI periods. F. Change in peak response amplitude (z-score) across training bins for neurons responsive with inhibition during early (0.5–1.2 s; left), mid (1.2–2.0 s; middle), and late (2.0–2.8 s; right) ISI periods. G. Heatmaps of single-neuron activity evoked by Cue A (left) and Cue C (right) sorted by the time of peak activity during trials 35–40 of Cue C pooled across mice. H. Correlation of time of peak activity of a given neuron evoked by each cue. Each point is a neuron colored by mice. Results are represented as mean ± SEM. ***p < 0.001; **p ≤ 0.01; *p ≤ 0.05.

